# Legacy effect of constant and diurnally oscillating temperatures on soil respiration and microbial community structure

**DOI:** 10.1101/2021.04.12.439414

**Authors:** Adetunji Alex Adekanmbi, Xin Shu, Yiran Zhou, Liz J. Shaw, Tom Sizmur

## Abstract

Laboratory incubation studies evaluating the temperature sensitivity of soil respiration often use measurements of respiration taken at a constant incubation temperature from soil that has been pre-incubated at the same constant temperature. However, such constant temperature incubations do not represent the field situation where soils undergo diurnal temperature oscillations. We investigated the effects of constant and diurnally oscillating temperatures on soil respiration and soil microbial community composition. A grassland soil from the UK was either incubated at a constant temperature of 5 °C, 10 °C, or 15 °C, or diurnally oscillated between 5 °C and 15 °C. Soil CO_2_ flux was measured by temporarily moving incubated soils from each of the abovementioned treatments to 5 °C, 10 °C or 15 °C, such that soils incubated at each temperature had CO_2_ flux measured at every temperature. We hypothesised that, irrespective of measurement temperature, CO_2_ emitted from the 5 °C to 15 °C oscillating incubation would be most similar to the soil incubated at 10 °C. The results showed that both incubation and measurement temperatures influence soil respiration. Incubating soil at a temperature oscillating between 5 °C and 15 °C resulted in significantly greater CO_2_ flux than constant incubations at 10 °C or 5 °C, but was not significantly different to the 15 °C incubation. The greater CO_2_ flux from soils incubated at 15 °C, or oscillating between 5 °C and 15 °C, coincided with a depletion of dissolved organic carbon and a shift in the phospholipid fatty acid profile of the soil microbial community, consistent with the thermal adaptation of microbial communities to higher temperatures. However, diurnal temperature oscillation did not significantly alter Q10. Our results suggest that daily maximum temperatures are more important than daily minimum or daily average temperatures when considering the response of soil respiration to warming.

## 2.0 Introduction

Soils harbour the largest actively cycling pool in the carbon cycle (Harden et al., 2018). Depletion of soil organic matter releases CO_2_, a greenhouse gas, into the atmosphere, contributing to global warming. The resulting increase in global temperatures is expected to stimulate heterotrophic soil respiration (Bardgett et al. 2008; Walker et al. 2018), thus causing a positive feedback that releases more CO_2_ into the atmosphere. The annual release of CO_2_ from soils by heterotrophic microorganisms is about 8 to 9 times higher than anthropogenic emissions from the burning of fossil fuels (Dutta and Dutta, 2016), so reducing the uncertainty concerning the magnitude of the positive feedback under future climate change scenarios deserves attention (Davidson and Janssens, 2006).

The impact of environmental change on soil carbon can be simulated using soil carbon models (e.g. ECOSSE (Dondini et al., 2016), DNDC (Gilhespy et al., 2014) and CENTURY (Parton, 1996)). These models divide the soil organic matter into pools which have different mean residence times in soil, related to their chemical recalcitrance. The rate of decomposition of each pool is subject to a first-order decay process, dependent, among other parameters, on the soil temperature, according to Arrhenius kinetics (Schimel and Weintraub, 2003). Other models (e.g. CEM (Foereid et al., 2014), CASA (Potter et al., 1993) and TEM (Raich et al., 1991) use a fixed Q_10_ value that represents the increase in soil respiration that occurs after a 10 °C increase in soil temperature (Meyer et al., 2018). These approaches have received criticism (Davidson and Janssens 2006; Schmidt et al. 2011) and, as a result, a new generation of soil carbon models are under development that better represent the physical and biological processes mediating soil organic matter turnover (Todd-Brown et al. 2012;Wieder et al. 2013;Abramoff et al. 2017). Soil carbon models generally operate on a monthly time step, using average monthly temperature as an input variable (Kirschbaum 1995; Yokozawa et al. 2010; Karhu et al. 2014), although there has been attempts to model daily time steps, using daily average temperature as an input variable (Gilhespy et al. 2014). In all of the abovementioned models, no consideration is made concerning the extent to which soil temperature oscillates diurnally, the influence this may have on the inherent temperature sensitivity of soil respiration, or whether daily or monthly average temperatures are adequate to capture the temperature sensitivity of soil respiration to the changes in temperature that we actually expect soils to experience (Mitra et al., 2019). There is a lack of experimental evidence highlighting the importance of diurnal temperature range, daily maximum temperature, or daily minimum temperature on soil respiration.

Assumptions on the relationship between soil temperature and soil respiration and their interpretation are often arbitrary (Subke and Bahn, 2010). One such facet that is arbitrary in nature is the selection of two temperatures for which the temperature coefficient (Q10) is determined during ex situ measurements (Graf et al., 2008). Often, the two temperatures, 10°C apart, that are chosen do not fall within the daily temperature ranges that soil microbial communities were previously exposed to in the field. To improve the accuracy with which temperature sensitivity (Q10) is determined in laboratory assays, estimating Q10 using temperature that reflect field conditions has been suggested (Pavelka et al., 2007; Graf et al., 2008). A similar diurnal oscillation of CO_2_ efflux and temperature (Akinremi et al., 1999) confirms the need for such recommendations. Estimating temperature sensitivity Q10 using the daily maxima and minima temperature may provide a better estimate of the relationship between soil temperature and soil respiration for modelling purposes.

The wide range of Q_10_ values reported in the literature may also be explained by differences in the laboratory and field procedures that have been used to measure soil respiration (Smith et al., 2018). A possible reason for this wide range may be because the temperature sensitivity of soil respiration is often assessed without considering that temperature oscillates diurnally (Ross and Täte, 1993;Winkler et al., 1996;Conant et al., 2008;Chen et al., 2009). This omission may be the reason why differences in seasonal Q10 of soil respiration earlier reported did not represent differences in the temperature sensitivity of soil microbial metabolism (Curiel Yuste et al., 2004;Gritsch et al., 2015). Periodic fluctuations in air temperatures can influence the soil temperatures and affect the underlying soil microbial activities (Chang et al., 2011) which, in turn, may influence soil respiration. How temperature fluctuations alter soil microbial community assemblages and how this may influence soil microbial function is important in predicting the impact of climate change on soil respiration (Uvarov et al. 2006; Hawkes and Keitt 2015).

Oscillating temperatures are uncommon in laboratory experiments, where soils are often incubated under constant temperatures for several months (von Lützow and Kögel-Knabner, 2009;Yan *et al*., 2017). Diurnal variation in the rates of soil microbial functions that release atmospheric gases such as CO_2_, N_2_O and CH_4_ have been reported in both the field (Zhou et al. 2015) and laboratory (Xu et al. 2016) studies. Such diurnal variation has been attributed largely to temperature oscillations (Shurpali et al., 2016). In addition to the abovementioned direct impacts of temperature on soil respiration that are due to increases in the activity of soil microbial communities, indirect effects on microbial activity could occur due to long term shifts in soil microbial community composition as a result of thermal adaptation (Luo et al. 2001; Davidson et al. 2006; Bradford et al. 2008; 2010; Buysse et al. 2013). Understanding how the legacy effects of changes in temperature regimes influence the structure and function behaviour of soil microbial communities is currently among the most important areas of investigation in the field of microbial ecology (Antwis et al. 2017). This area is important because, along with a future increase in global mean temperatures, we also expect a dampening of the diurnal temperature range, with daily minimum temperatures expected to increase more than daily maximum temperatures (Braganza et al. 2004; Zhou et al. 2009). Thus, soil microorganisms may become thermally adapted to a narrower diurnal temperature range and respond differently to temperature increases than the communities that currently inhabit soils.

Measuring soil respiration in soils that are incubated at diurnally oscillated temperature may create conditions that are more similar to those experienced by soil microbial communities in nature (Thiessen et al., 2013). However, previous attempts at measuring soil respiration under controlled oscillating temperatures are rare. A few studies have achieved this simulated oscillation by moving soils from one temperature to another and holding them at these constant temperatures for longer than may occur in nature (e.g. between 9 and 12 hours), during which soil respiration measurements were made (e.g. Fang et al. 2005;Thiessen et al. 2013). Unfortunately, these studies did not report how microbial community composition changed as a result of these oscillating soil temperatures. Past temperature regimes already experienced by soil can influence both the soil microbial community (Wu et al., 2010) and substrate availability through depletion (Pold et al., 2017). It is uncertain how current respiration and temperature sensitivity of respiration depends on the interaction between changes in soil microbial community and substrate depletion due to the legacy effect of the past temperature regime already experienced by the soil. Comparing the soil microbial community composition and its function *ex situ* under both constant and diurnally oscillating temperatures that mimic real diurnal temperature oscillations may offer a better understanding of how soil microbial communities may change under future environmental change and help us to better predict the magnitude of the positive feedback of CO_2_ flux into the atmosphere.

We designed and executed a laboratory incubation experiment to examine the effects of constant and diurnally oscillating temperatures on soil microbial community structure and function. The temperature treatments were chosen to reflect average daily minimum, daily average, and daily maximum temperatures in Reading, UK. Soils were incubated at these three constant temperatures (5 °C, 10 °C or 15 °C) alongside soils that were oscillated between daily minimum (5 °C) and daily maximum (15 °C) temperatures. Soil CO_2_ flux was measured by temporarily moving incubated soils from the abovementioned treatments to 5 °C, 10 °C or 15 °C, such that soils incubated at each temperature had CO_2_ flux measured at every temperature. Our approach used incubation and measurement temperatures as statistical factors to explore the influence of incubation temperature on the respiration at the measured temperature. Our aim was to determine whether soil samples incubated under different temperature regimes exhibit different respiration rates, even if the measurements of respiration are all made at the same temperature. This approach availed us the opportunity to calculate Q10 using the 5 °C and 15 °C measurement temperatures. We hypothesised that 1) respiration measurements made at higher temperatures would result in greater respiration rates, 2) incubation temperature will induce changes in soil microbial community structure and the availability of soil C and N, and that 3) these changes in community composition and biogeochemistry would lead to different respiration rates from soils incubated under different temperature regimes, even when respiration was measured at the same temperature. Our null hypothesis was that soil respiration of soils diurnally oscillating between 5 °C and 15 °C would be similar to respiration from soils incubated at 10 °C, thus confirming the suitability of daily or monthly average temperatures as input variables in soil carbon models.

## 3 Materials and Methods

### 3.1 Site selection and Soil Sampling

Soil was collected at 0-10 cm depth from a permanent grassland field (Latitude 51°28.564′, Longitude 000 °54.198′) on the University of Reading experimental farm at Sonning, UK. The soil was identified as a Chromic Endoskeletic Luvisol. Details of the soil description and land use history are provided in Adekanmbi et al. (2020). Multiple subsamples from an area of approximately 10 m^2^ were bulked together to obtain a composite sample. The fresh soil was sieved to 4 mm, thoroughly mixed, and then stored at 4°C until the start of the experiment. A subsample of approximately 500 g was air-dried to characterise soil texture, water holding capacity (%WHC), pH in water, total carbon (TC), total nitrogen (TN), and available NO_3_^-^ and NH_4_^+^ (Table 1). The methods for each of these analyses are reported in the supplementary material.

**Table 1.**
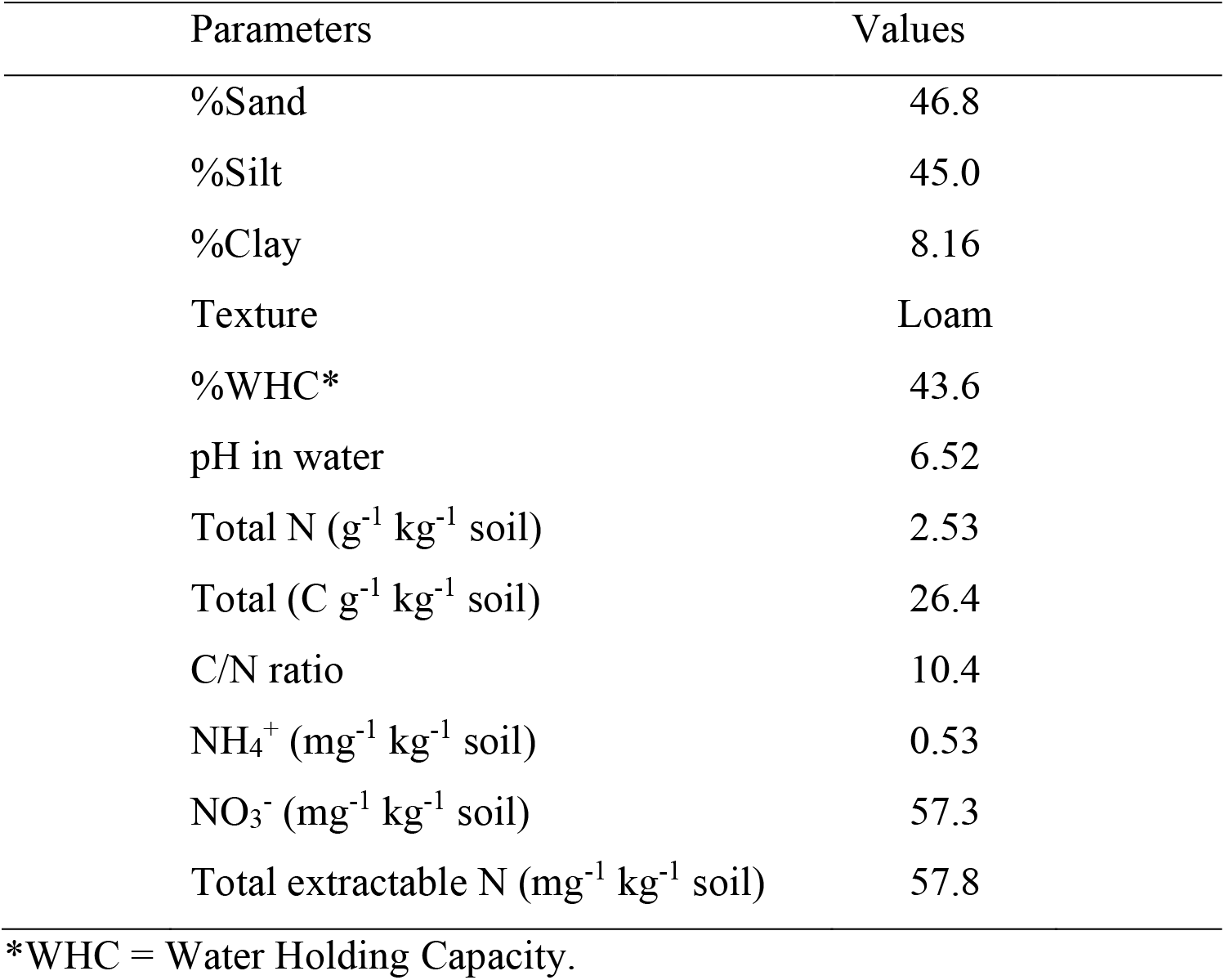
Physical and chemical properties of the soil used in the experiment.

### 3.2 Experimental Design

The experiment was a 4 x 3 factorial design comprising of 4 incubation temperatures (5 °C, 10 °C, 15 °C, or diurnally oscillating between 5 °C and 15 °C) and 3 measurement temperatures (5°C, 10 °C, and 15 °C), with four replicates (Table 2), resulting in 12 treatments and 48 units (Treatments 1 – 12 in Table 2). Each week of the experiment the soil samples were incubated in controlled environment chambers for six days at their allocated incubation temperatures before moving to their allocated measurement temperatures 24 hours prior to respiration measurement, and then returned to their allocated incubation temperature after measurement of CO_2_ flux (Figure 1A and 1B). Two blank (without soil) incubation jars were incubated at each measurement temperature as a blank to correct for background atmospheric CO_2_ concentration in the mesocosms and accounted for while calculating the CO_2_ flux.

**Table 2.** **Experimental design highlighting the incubation and measurement temperatures and the analysis undertaken on experimental units assigned to individual treatments.**

| Treatment Number | Incubation Temperature | Measurement Temperature | Replicates | CO<sub>2</sub> flux | Post incubation analysis |
| --- | --- | --- | --- | --- | --- |
| 1 | 5°C | 5°C | 4 | ✓ | ✓ |
| 2 | 5°C | 10°C | 4 | ✓ |  |
| 3 | 5°C | 15°C | 4 | ✓ |  |
| 4 | 10°C | 5°C | 4 | ✓ |  |
| 5 | 10°C | 10°C | 4 | ✓ | ✓ |
| 6 | 10°C | 15°C | 4 | ✓ |  |
| 7 | 15°C | 5°C | 4 | ✓ |  |
| 8 | 15°C | 10°C | 4 | ✓ |  |
| 9 | 15°C | 15°C | 4 | ✓ | ✓ |
| 10 | Oscillating<br>(5°C -15°C) | 5°C | 4 | ✓ |  |
| 11 | Oscillating<br>(5°C -15°C) | 10°C | 4 | ✓ |  |
| 12 | Oscillating<br>(5°C -15°C) | 15°C | 4 | ✓ |  |
| *13 | Oscillating<br>(5°C -15°C) | Oscillating<br>(5°C -15°C) | 4 | ✓ | ✓ |

\* Treatment 13 was not included in statistical analysis for CO<sub>2</sub> flux

**Figure 1.**
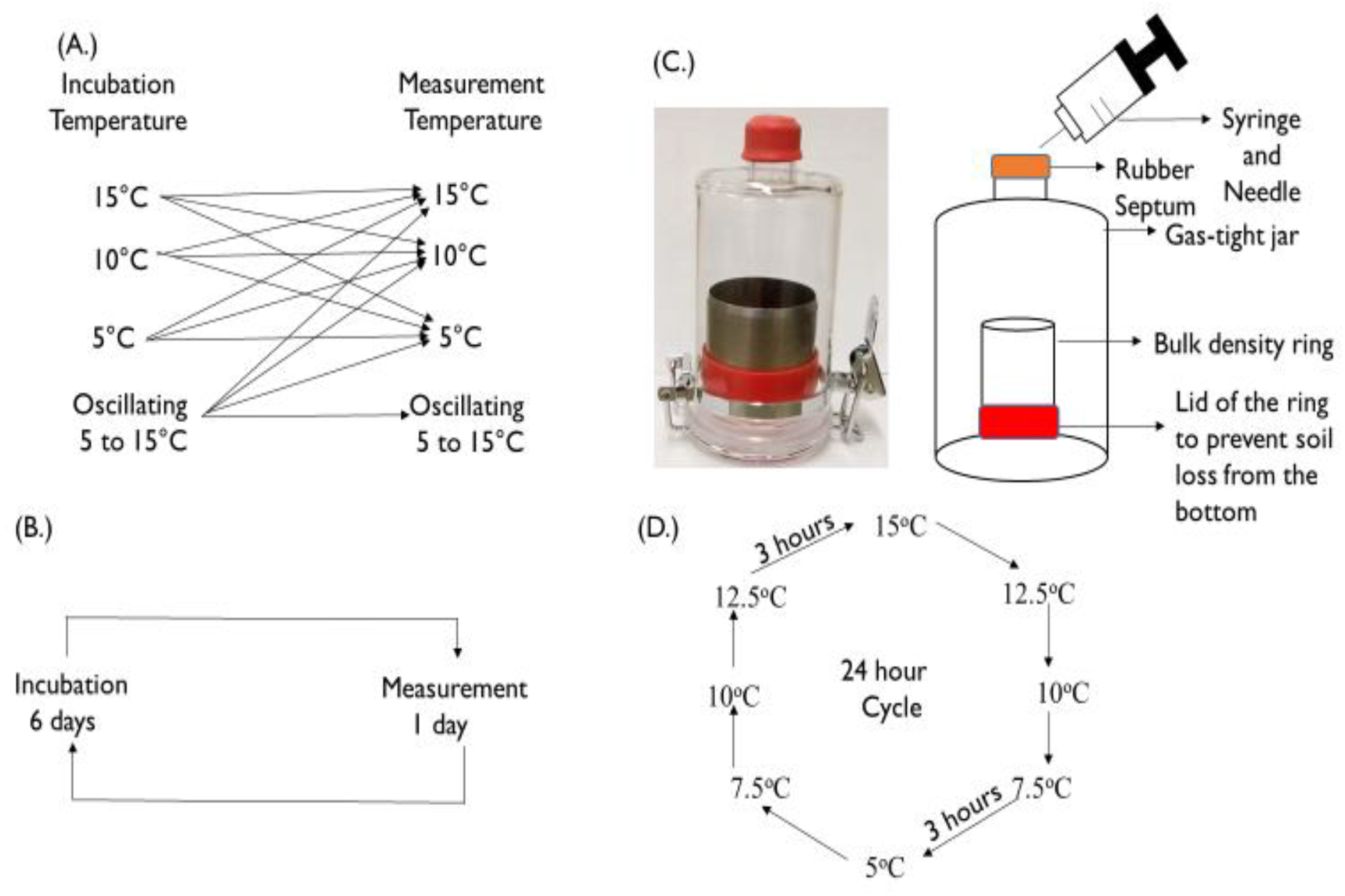
Experimental design including (A) a graphical depiction of the experimental treatments showing how individual treatments were moved from their incubation temperature to their measurement temperature prior to CO_2_ flux measurements; (B) the weekly schedule for moving soils from their incubation temperatures to their measurement temperatures; (C) the design of the incubation containers and the method by which headspace gas samples were collected from the soil incubation containers; and (D) the daily temperature regime that the soils assigned to the 5°C to 15°C oscillating treatment were exposed to, with the temperature held for three hours at each temperature step.

Four extra cores were both incubated and measured in an environment diurnally oscillating between 5 °C and 15 °C (See treatment 13 in Table 2). Measurements of CO_2_ flux were made when the environment was at 10 °C while the temperature was decreasing during the diurnal oscillation. The addition of this treatment meant that, at the end of the experiment, we had soils that had remained (without movement) at 5 °C, 10 °C, 15 °C, and diurnally oscillating between 5 °C and 15 °C (representing 4 treatments, and 16 experimental units). These units were used for post incubation soil chemical and biological analysis.

### 3.3 Experimental setup and CO_2_ flux measurements

Field moist soil samples of 70 g fresh weight (equivalent to 56.51 g dry weight) were weighed into a 5 x 5 cm cylinder (height x diameter; volume = 98.22 cm^3^) and placed in a 320 ml gas-tight container (Figure 1C). The containers were modified to allow gas collection ports, which were covered with Parafilm® to reduce moisture loss (but allow gas exchange) when not in use, following Adekanmbi et al. (2020). The soils were adjusted to 60% of their water holding capacity, as described by Yang et al., (2017). All the soil samples were pre-incubated for 7 days at their respective incubation temperature to allow the sieving/re-wetting induced flush in respiration (Liu et al., 2018) to subside before the first CO_2_ flux measurement was made. The temperatures selected for our experiment were the average daily minimum (5 °C), average daily maximum (15 °C), and average daily mean (10 °C) temperatures measured over a 28 year (1 January 1990 – May 2018) period at the University of Reading Meteorological station, situated approximately 2.5 miles from University of Reading experimental farm at Sonning, where the soil for this experiment was collected. We set the oscillating treatment to oscillate diurnally between the average daily minimum (5 °C) and average daily maximum (15 °C) by programming a growth chamber to spend three hours at each of eight temperatures per day (5 °C, to 7.5 °C, to 10 °C, to 12.5 °C, to 15 °C, to 12.5 °C, to 10 °C, to 7.5 °C, and then back to 5 °C), as shown in Figure 1D.

The experiment lasted for 119 days (17 weeks). Soil respiration was measured as CO_2_ flux every week up until the third week (day 21) and then at two-week intervals thereafter until the 17^th^ week. Prior to each CO_2_ flux measurement, the Parafilm® was removed to allow the gas in the gas-tight container to mix with the atmosphere. During CO_2_ flux measurement, containers were sealed with a Suba-Seal® Septa and kept at the measurement temperature for one hour before a 15 ml headspace gas sample was taken from each container using a syringe and hypodermic needle and transferred into a pre-evacuated Labco® exetainer vial. After each sampling, the septum was removed and the Parafilm® replaced to reduce moisture loss. The gas samples were analysed using an Agilent 7890A (Agilent Technologies, Wilmington USA) gas chromatograph fitted with a 1/8 inch stainless steel packed column (HayeSep Q 80/100) to separate the CO_2_ peak in an oven held at 60 °C using N_2_ as the carrier gas at a rate of 21 ml min^-1^ prior to conversion to CH_4_ in a methanizer and detection using a flame ionization detector. The moisture content of the soil in each container was adjusted back to 60% of their water holding capacity after collecting gas and before returning samples back to their incubation temperatures by addition of deionised water to compensate for mass loss due to evaporation.

### 3.4 Laboratory analysis of soil chemical and biological properties

At the end of the experiment (after 17 weeks), soil samples were taken from the 16 containers that had remained (without movement) at 5 °C, 10 °C, 15 °C, or diurnally oscillating between 5 °C and 15 °C for the entirety of the experiment to examine soil chemical and biological properties. A 10 g sub-sample of soil was extracted immediately for determination of available NO_3_^-^ and NH_4_^+^. A further 5 g was used to determine the gravimetric water content and adjust the results of the NO_3_^-^ and NH_4_^+^ analysis for soil moisture so that they could be expressed on a dry mass basis. A 5 g sub-sample was freeze-dried prior to phospholipid fatty acid (PLFA) analysis. A 15 g sub-sample was air-dried for chemical analysis to determine TC, TN, and hot and cold water extractable carbon (HWEOC and CWEOC).

Soil microbial community structure was assessed using PLFA profiles (Tunlid and White, 1992). Freeze-dried soils (2 g per sample) were extracted using Bligh and Dyer solvent (Bligh and Dyer, 1959) following Frostegård and Bååth (1996). Extracted phospholipids were derivatised, as described by Dowling *et al*. (1986), and analysed as fatty acid methyl esters by gas chromatography (Agilent 6890N, flame ionization detector and a 30 m x 0.25 mm capillary column with a 0.25 μm film of 5% diphenyl, 95% dimethyl siloxane) following Frostegård *et al*. (1991). Individual fatty acid methyl esters were identified and quantified according to the retention times and peak area in using quantitative and qualitative standards (26 bacterial FAMEs, C11 to C20 and 37 FAMEs, C4 to C24; Supelco, Supelco UK, Poole, UK). Individual PLFAs were attributed to microbial groups according to (Kaur et al., 2005; Willers et al., 2015; and Quideau et al., 2016). The assignment of individual fatty acid biomarkers to microbial groups or stress indices is described in Table S-1 in the supplementary material.

Total C and N were analysed using a C/N elemental analyser (Thermo Scientific Flash 2000 EA). Available NO_3_^-^ and NH_4_^+^ were analysed by extracting 10 g moist soil with 50 ml of 1M KCl for 1 hour, filtering (GF/A 15.0cm diameter), and analysing using a Skalar SAN++ flow injection auto-analyser. HWEOC and CWEOC were analysed as described by Ghani, et al., (2003). Approximately 3 g air-dried soil samples of known moisture content were accurately weighed into 50 ml polypropylene centrifuge tubes. Thirty ml of ultra-pure water was then added to each tube before mixing on a rotary shaker at 30 rpm for 30 min at 20°C. This was followed by centrifuging at 3500 rpm for 20 min at 20 °C. The supernatants were then removed using polypropylene syringes and passed through 0.2 µm cellulose nitrate membrane filters into polypropylene universal tubes, discarding the first 3 ml of the filtrate each time. A further 30 ml of ultra-pure water was then added to each centrifuge tube before vortexing for 10 seconds and leaving in an 80 °C water bath overnight and the supernatants removed, as described above. Both supernatants were analysed for CWEOC and HWEOC, respectively using a Shimazu TOC analyser.

### 3.5 Q10 Determination

The temperature sensitivity was determined by calculating the temperature coefficient (Q10) using the equal time method (Lin et al., 2015; Zang et al., 2020):

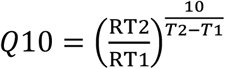

where RT2 is CO_2_ flux measured at 15 °C, RT1 is the corresponding CO_2_ flux measured at 5 °C, T2 is 15 °C, and T1 is 5 °C.

### 3.6 Statistical Analysis

Differences in soil respiration due to incubation and measurement temperatures over the period of 17 weeks were tested using repeated measures analysis of variance (ANOVA) and Fisher’s Least Significant Difference (LSD) test was used for pairwise comparisons in Genstat (10^th^ edition). Two way ANOVA was conducted to assess effect of incubation temperature and incubation week on Q10. Non-metric multi-dimensional scaling (NMDS) ordination derived from Bray-Curtis similarities was used to separate soil microbial community structures of samples subjected to different incubation temperatures using the *vegan* package (Oksanen, 2017). The distance was Bray Curtis, performed in 2 dimensions, with stress factor of 0.05370305. PERMANOVA was used to assess whether incubation temperature influenced the soil microbial community distance. R v3.5.1 (R Development Core Team, 2018) was used to perform the NMDS. One-way ANOVA was also used to assess the differences in the NMDS 1 and 2 after establishing that there was a significant difference in community distance due to temperature to examine the direction of temperature impact on soil microbial community. One-way ANOVA was used to test the differences in soil properties and PLFA biomarkers due to incubation temperatures.

## 4 Results

### 4.1 Effects of measurement and incubation temperatures on soil CO_2_ flux

The CO_2_ flux data for each individual treatment are presented in Figure S-1 and Figure S-2 of the Supplementary material. Repeated measures ANOVA of the data revealed that both incubation (*P* < 0.001) and measurement (*P* < 0.001) temperature had a significant effect on the CO_2_ flux measured (Table 3). However, there was no significant interaction between incubation and measurement temperatures (*P* = 0.680). Irrespective of measurement temperature, incubating soil at 15 °C, or oscillating between 5 °C and 15 °C, released significantly (*P* < 0.001) more CO_2_ than incubating at 5 °C or 10 °C (Figure 2A, Table 3). As expected, CO_2_ flux, when measured at 15 °C, was significantly (*P* < 0.001) greater than CO_2_ flux measured at 5 °C or 10 °C (Figure 2B, Table 3). However, counter to expectations, it was observed that soils measured at 5 °C released slightly (but not significantly) more CO_2_ compared to soils incubated at 10 °C (Figure 2B). Irrespective of measurement temperature, CO_2_ flux from the soils that oscillated between 5 °C and 15 °C was not significantly different (*P* > 0.05) from the soils incubated at 15 °C, but was significantly (*P* < 0.001) greater than the soils incubated at 10 °C or 5 °C (Figure 2A).

**Table 3.**
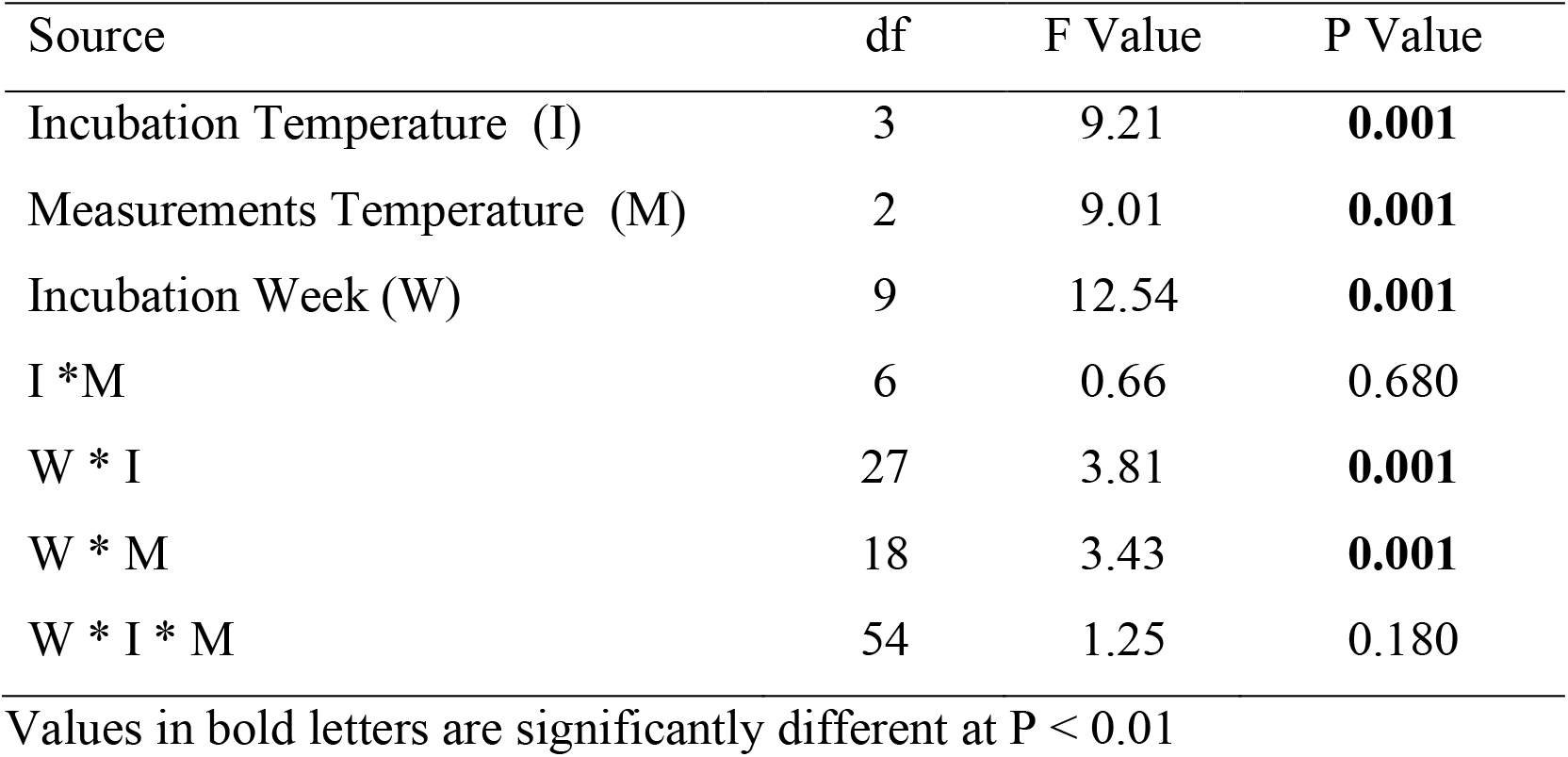
**Summary table for two-way repeated measures ANOVA for soil respiration; Incubation and Measurement temperatures were the main (subject) factors.**

**Figure 2:**
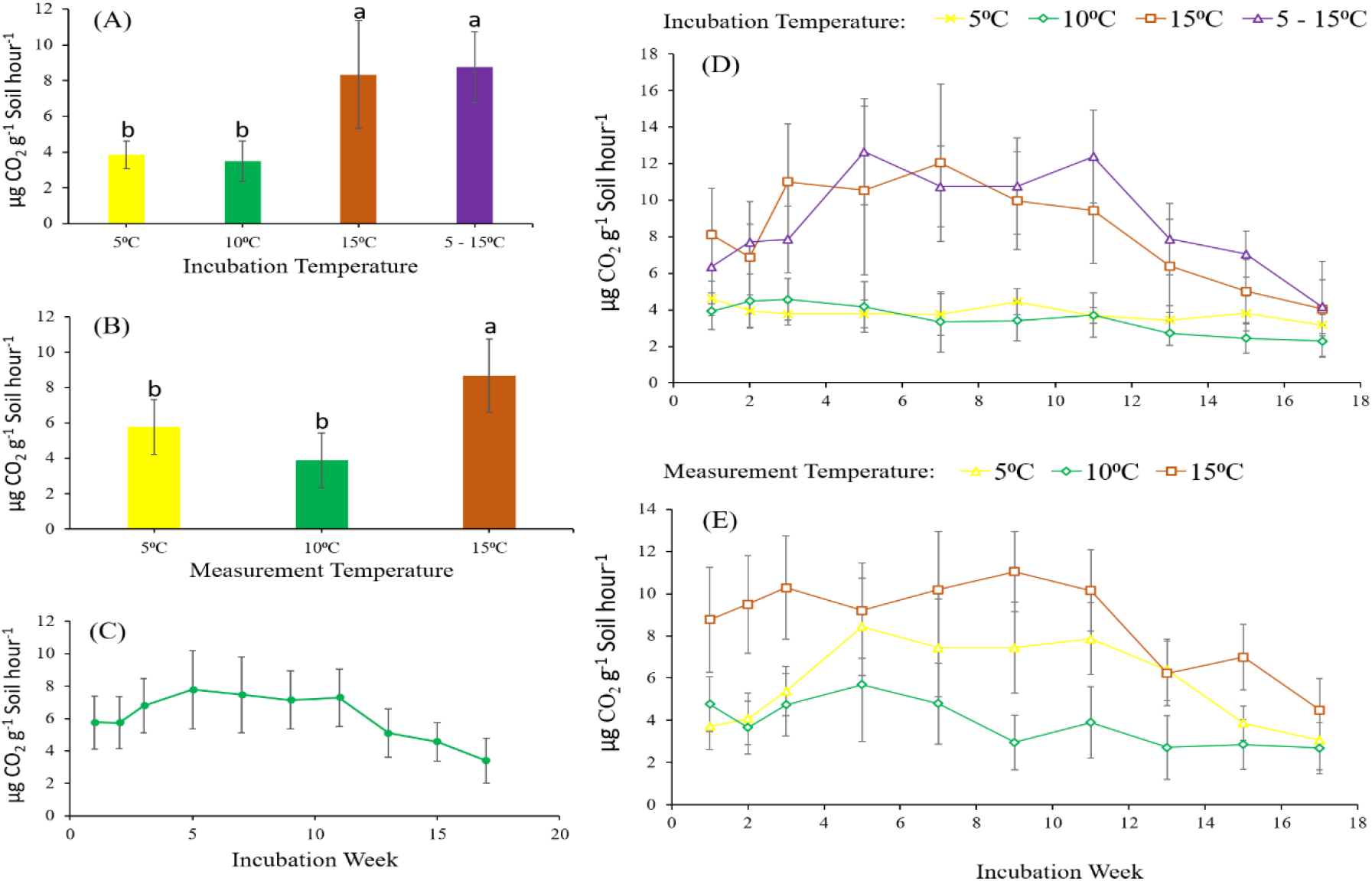
**Effects of soil incubation temperature (A), measurement temperature (B), Incubation Week (C), Incubation Week x temperature (D) and Incubation Week x Measurement temperatures (E) on soil CO_2_ flux from soil. Error bars represent standard errors of the mean. Bars with the same lower case letters are not significantly different from each other (*P* > 0.05). For (A) n = 120, for (B) n = 160, for (C) n = 48, for (D) n = 12, and for (E) n = 16.**

Repeated measures ANOVA of the data also revealed that incubation week (*P* < 0.001) had a significant effect on the CO_2_ flux measured (Table 3), as indicated by a slightly elevated CO_2_ flux measured between week three and week eleven (Figure 2C). There was also a significant interaction (*P* < 0.001) between incubation week and incubation temperature, and between incubation week and measurement temperature on soil CO_2_ flux (Figure 2D and 2E, Table 3). The abovementioned elevated CO_2_ flux measured between week three and eleven was more pronounced in treatments incubated at 15 °C or oscillating between 5 °C and 15 °C (Figure 2D). Although CO_2_ flux measured at 5 °C was lower than that measured at 10 °C in week 1 of the experiment, it was greater in soils measured at 5 °C for the remainder of the experiment (Figure 2E).

### 4.2 Effects of incubation temperature on temperature sensitivity (Q10) of Soil respiration

The result on the effects of incubation temperature on the Q10 of soil respiration is presented in Figure 3. The two-way analysis of variance (data not shown) showed that, incubation temperature (*P* < 0.0001), but not incubation week (*P* = 0.504), significantly influenced the calculated Q10. Irrespective of incubation week, incubating soil at 5 °C led to a significantly higher Q10 compared to soil incubated at 10 °C. Constantly incubating soil at 15 °C or at temperature oscillating between 5 and 15 °C had similar Q10 and this showed an intermediate Q10 between soil incubated at either 5 °C or 10 °C. The Q10 derived from incubating soil diurnally oscillating between 5 and 15 °C was not significantly different from that obtained using soils incubated constantly at 5, 10, or 15 °C.

**Figure 3:**
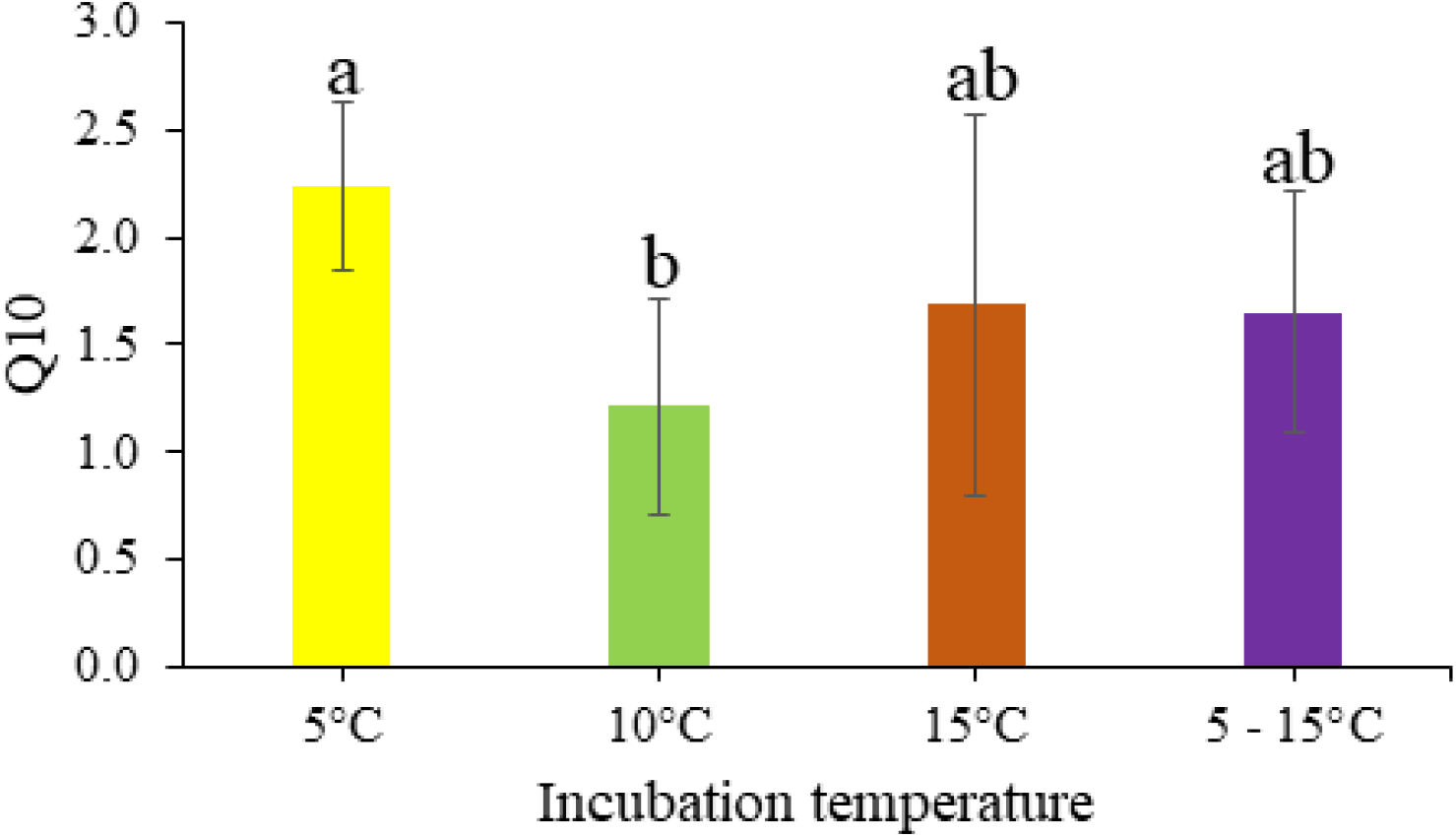
**Temperature sensitivity (Q10) of soil respiration as affected by incubation temperature. Bars and error bars are the means and standard error of data recorded weekly for 17 weeks. Bars that share the same letter within a graph are not significantly different from each other (*P* > 0.05).**

### 4.3 Effects of incubation temperature on soil carbon and nitrogen

The concentrations of chemical fractionations of C and N in soils from the 16 experimental units that remained (without movement) at the same temperature for the duration of the experiment (Table 2) are presented in Figure 4. CWEOC (Figure 4A; Table 4), was significantly (*P* < 0.05) higher in soils incubated at 5 °C and 10 °C compared to those in incubated at 15 °C or oscillated between 5 °C and 15 °C. Furthermore, soils incubated at 5 °C had a significantly (*P* < 0.05) higher HWEOC than all other incubation temperatures (Figure 4B; Table 4). TC (Figure 4C; Table 4) was significantly (*P* < 0.05) higher in soils incubated at 5 °C or 10 °C, compared to those incubated at 15 °C or oscillated between 5 °C and 15 °C. Total extractable N (TEN) significantly (*P* < 0.05) increased with incubation temperature (Figure 4D; Table 4). Soils oscillated between 5 °C and 15 °C had a similar TEN to soils incubated 15 °C, but significantly (*P* < 0.05) greater than soils incubated at 5 °C or 10 °C. The TN concentration was significantly (*P* < 0.05) greater in soils oscillated between 5 °C and 15 °C, compared to all other treatments incubated at constant temperatures (Figure 4E; Table 4). 63The C/N ratio significantly (*P* < 0.05) decreased with increasing incubation temperature, but, unlike TEN, soils oscillated between 5 °C and 15 °C had a significantly (*P* < 0.05) different (lower) C/N ratio than soils incubated at 15 °C (Figure 4F; Table 4).

**Table 4:** **Summary of ANOVA on the impact of temperature on Carbon and Nitrogen**

| Sources | d<br>f | F Values | P Values |
| --- | --- | --- | --- |
| CWEOC $\mu\text{g C g}^{-1}$ Soil | 3 | 9.38 | **0.002** |
| HWEOC $\mu\text{g C g}^{-1}$ Soil | 3 | 4.33 | **0.028** |
| TC $\text{g kg}^{-1}$ Soil | 3 | 5.55 | **0.013** |
| TEN $\text{mg kg}^{-1}$ Soil | 3 | 45.53 | **0.0001** |
| TN $\text{g kg}^{-1}$ Soil | 3 | 11.24 | **0.001** |
| C/N | 3 | 53.56 | **0.0001** |

Values in bold letters are significantly different at $P < 0.05$ ; C/H-WEOC = Cold/Hot water extractable carbon; TEN = Total extractable N.

**Figure 4:**
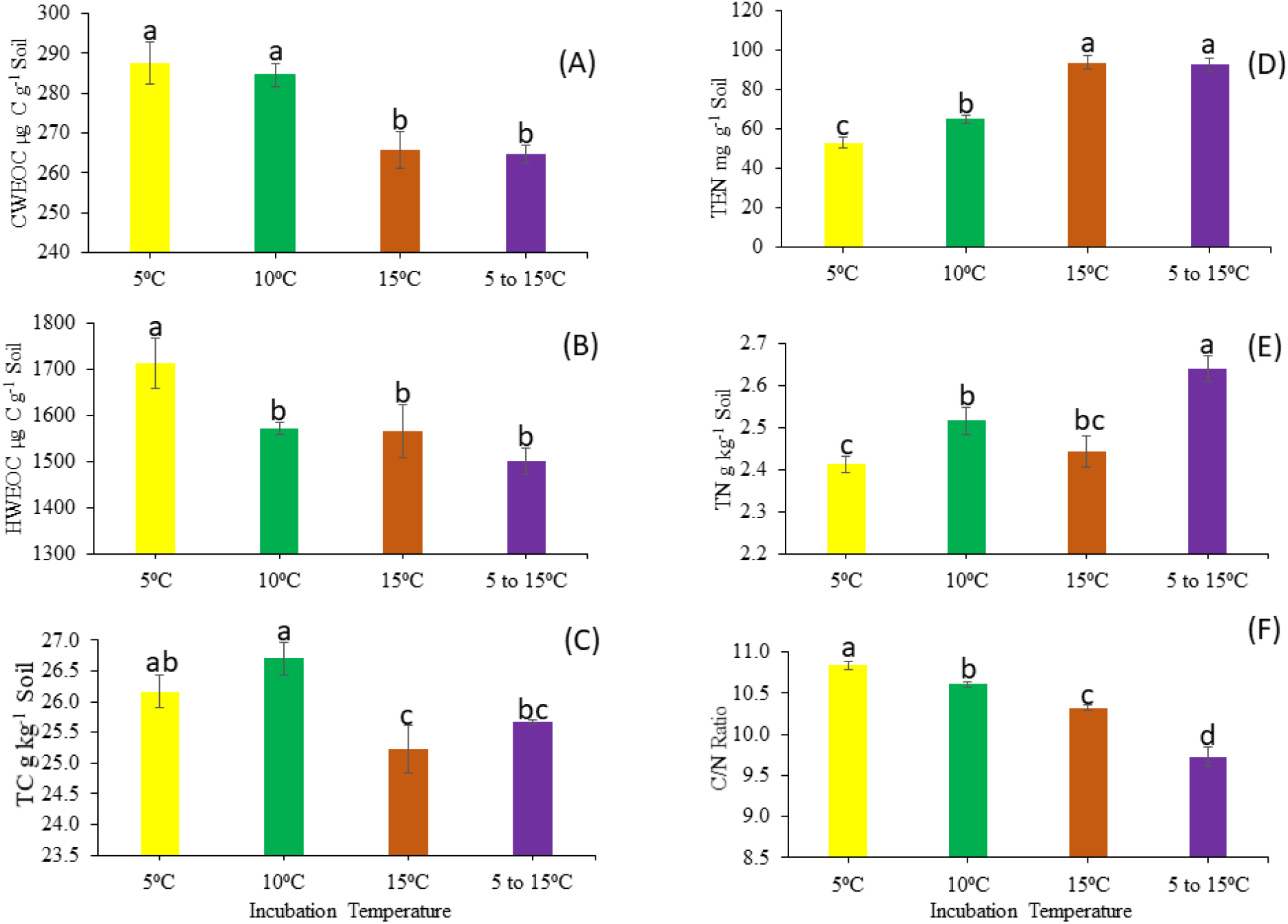
**Influence of incubation temperature on cold (CWEOC = A), and hot (HWEOC = B) water extractable carbon, Total Carbon (TC = C), Total extractable N (NH_4_^+^ and NO_3_^-^) (TEN = D), Total Nitrogen (TN = E), and C/N Ratio (F). Bars and error bars represent mean and standard error of the mean (n = 4). Bars that share the same letter within a graph are not significantly different from each other (*P* > 0.05).**

### 4.4 Impacts of incubation temperature on soil microbial community composition

The structure of the soil microbial community, as measured using PLFA biomarkers, was affected by soil incubation temperature, as shown in Figure 5. PERMANOVA analysis revealed that temperature had a significant (*P* = 0.002) effect on microbial community distance (Bray Curtis distance between the PLFA profiles). One-way ANOVA of NMDS score 1 (NMDS1) showed that the soil microbial community structure in soils incubated at 5 °C and 10 °C were not statistically different to one another, but distinct (*P* = 0.0001) from those in soils incubated at 15 °C, or oscillated between 5 °C and 15 °C (Figure 6, Table 5). There was a slightly (non-significant; *P* > 0.05) lower abundance of bacteria, and total PLFA (i.e. total microbial biomass) in soils incubated at 15 °C, or oscillated between 5 °C and 15 °C, compared to soils incubated at 5 °C or 10 °C (Figure 6A and 6B). The abundance of fungal biomarkers and the fungal/bacterial ratio was significantly (*P* < 0.05) lower in soils incubated at 15 °C or oscillated between 5 °C and 15 °C, compared to soils incubated at 5 °C or 10 °C (Figure 6D and 6E). Furthermore, the ratio of Gram-negative/Gram-positive bacteria was significantly (*P* < 0.05) greater in soils incubated at 5 or 10 °C compared to soils incubated at 15 °C or oscillating between 5 °C and 15 °C (Figure 6C; Table 5). Likewise, the ratios of (i) cis to trans isomers, (ii) iso to anteiso branching, and (iii) cyclopropyl fatty acids to their monoenoic precursors were all significantly (*P* < 0.05) higher in soils incubated at 15 °C or oscillating between 5 °C and 15 °C, compared to those incubated at 5 °C or 10 °C (Figure 6F, 6G and 6H; Table 5).

**Table 5:** **Summary ANOVA table for a one-way ANOVA on the impact of incubation temperature on the soil microbial community composition assessed using PLFA biomarkers**

| Sources | df | F Values | P Values |
| --- | --- | --- | --- |
| Total PLFA (nmol g<sup>-1</sup> soil) | 3 | 2.42 | 0.117 |
| Bacterial (nmol g<sup>-1</sup> soil) | 3 | 2.38 | **0.121** |
| G-/G+ Bacteria Ratio | 3 | 5.11 | **0.017** |
| Fungal (nmol g<sup>-1</sup> soil) | 3 | 5.51 | **0.013** |
| Fungal/Bacterial Ratio | 3 | 27.39 | **0.0001** |
| cis/trans Ratio | 3 | 18.36 | **0.0001** |
| iso/anteiso ratio | 3 | 31.87 | **0.0001** |
| cy17:0c/16:1ω7c | 3 | 58.32 | **0.0001** |
| PERMANOVA | 3 | 9.14 | 0.002 |
| NMDS1 | 3 | 36.85 | 0.0001 |
| NMDS2 | 3 | 0.52 | 0.675 |

Values in bold letters are statistically significant at $P < 0.05$

**Figure 5:**
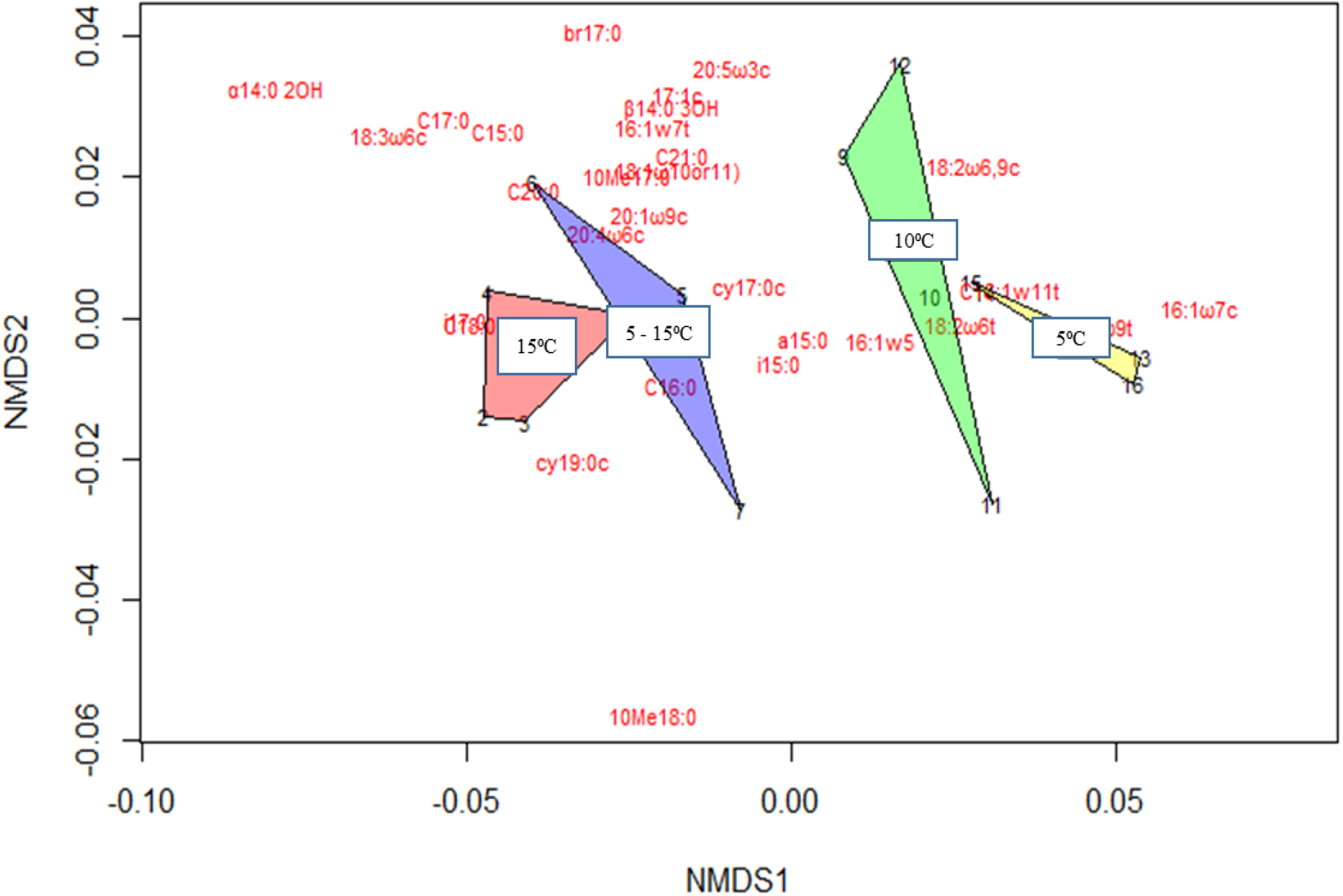
**NMDS plot showing the distribution of lipid biomarkers as influenced by soil incubation temperature on the soil microbial community structure measured using Phospholipid Fatty Acid Analysis. The distance is Bray Curtis, performed in 2 dimensions, with stress factor of 0.054. Shaded coloured regions encompass samples from the different incubation treatments (n = 4).**

**Figure 6:**
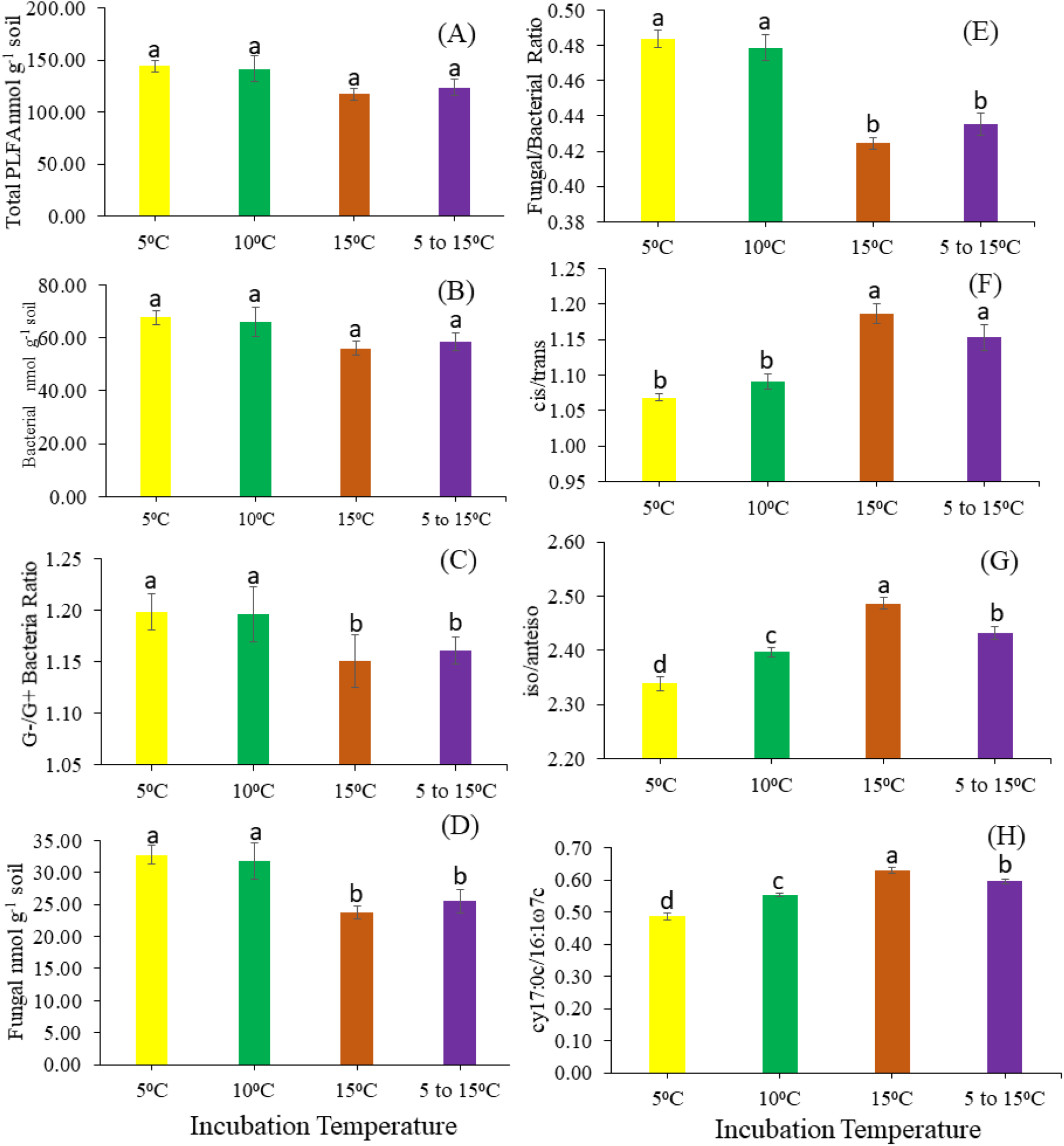
**Influence of incubation temperature on total PLFA (A), bacterial abundance (B), gram-negative/gram-positive bacterial ratio (G-/G+) (C), fungal abundance (D), fungal/bacterial ratio (E) cis/trans isomer ratio (F), iso/anteiso braching ratio (G) and cyclpropyl to monoenoic precursor (cy17:0c/16:1ω7c) ratio (H). Bars and error bars represent mean and standard error of the mean (n = 4). Bars that share a letter are not significantly different from one another (*P* < 0.05).**

## 5 Discussion

### 5.1 The legacy of previous incubation temperature on soil respiration

It is well established in multiple soil warming experiments undertaken in the field and laboratory that soil respiration increases in response to temperature rises (Chen et al. 2000; von Lützow and Kögel-Knabner 2009; Bell et al. 2010; Karhu et al. 2014; Carey et al. 2016; Melillo et al. 2017). Faster metabolism of microbially available organic carbon is the major reason suggested for the increases in soil CO_2_ flux observed (Zogg et al. 1997; Melillo et al. 2017; Walker et al. 2018). Counter to expectations, the incubation temperatures had a larger impact on CO_2_ flux than the temperature at which CO_2_ flux was measured (Figure 2 and Table 3) whereby CO_2_ flux was greater from soils that had been incubated at higher incubation temperatures, regardless of measurement temperature. Our observation implies that the temperature that a soil has previously been exposed to can exert a considerable legacy effect on the future soil respiration rate.

A possible explanation for this observation may be related to the knowledge that extracellular depolymerisation of macromolecular organic carbon is considered the rate-limiting step in the mineralisation of soil organic matter and that soil microorganisms invest more into extracellular enzyme excretion when placed under resource limited conditions (Allison, 2014). Jan et al. (2009) demonstrated that protein mineralisation to CO_2_ is 20 times slower than amino acid mineralisation to CO_2_ and is highly temperature sensitive. In soils incubated at 15 °C, or oscillating between 5 °C and 15 °C, the soil microbial community may have thermally adapted to produce a greater quantity of extracellular enzymes that depolymerise recalcitrant substrates (Meng et al., 2020), due to labile substrate depletion. The abundance of these extracellular enzymes in the soil environment during the measurement of respiration may have been adequate to depolymerise sufficient macromolecules to prevent the availability of low molecular weight compounds from being the rate limiting factor mediating respiration at any CO_2_ flux measurement temperature. Extracellular enzymes could continue to depolymerise even at low measurement temperatures when microbial uptake of the produced monomers ceases due to temporary reductions in membrane fluidity (Nedwell, 1999). Concurrently, in soils incubated at 5 °C or 10 °C, lower extracellular enzyme activity may have resulted in lower availability of low molecular weight compounds and thus a legacy of previous incubation temperature regime on soil respiration. This finding has important implications for soil scientists who pre-incubate soils prior to making respiration measurements. Broadly speaking, pre-incubation at high temperatures may result in substrate depletion and greater production of C-acquiring extracellular enzymes, increasing the probability that intracellular respiration becomes the rate limiting step during the measurement of respiration. Conversely, pre-incubation at low temperature may result in extracellular depolymerisation being the rate limiting step in respiration. Unfortunately, neither of these circumstances reflect the real diurnal oscillations that soils experience in the field.

We observed similar CO_2_ flux from soils that were incubated constantly at 15 °C and soils incubated at diurnally oscillating soil temperature between 5 °C and 15 °C (Figure 2A). We thus conclude that the time spent at 15 °C in the diurnally oscillating treatment was sufficient for soil microbial communities to produce extracellular enzymes to depolymerise enough macromolecules to prevent the availability of low molecular weight compounds from being the factor limiting the rate of intracellular respiration. This result also implies that maximum daily temperature is an important factor influencing the transformation of soil organic carbon to CO_2_; perhaps more important than daily average temperature. This assertion has important implications for our predictions of the effect that future environmental change may have on the global carbon cycle. The last half of the 20th century saw daily minimum temperatures increased by 0.9 °C while daily maximum temperatures increased by only 0.6 °C (Braganza et al., 2004). Therefore, while the climate warms, we are experiencing a reduction in the diurnal temperature range (Alexander et al., 2006) due to increased cloud cover and sulphate aerosol emission (Hansen et al., 1995). It is thus imperative to ensure that the next generation of land-surface models adequately simulate the impact of this asymmetric warming on the production and activity of extracellular enzymes and the subsequent impacts on soil heterotrophic respiration.

We also observed that the temperature a soil had been previously incubated at influenced the Q10 calculated from the measurement of CO_2_ flux at 5 °C and 15 °C (Figure 3). Soil previously incubated at 5 °C resulted in the highest Q10 and soil previously incubated at 10°C resulted in the lowest Q10. It is commonplace to refrigerate soils after field collection to supress microbial activity prior to making respiration measurements used to generate Q10 values(Gritsch et al., 2015;Li et al., 2015;Meyer et al., 2019). This activity may result in an overestimation of the Q10 because pre-existing extracellular enzymes may still be able to depolymerise macromolecules and generate a pool of labile carbon (Figure 4) that is not mineralised by microorganisms due to low membrane fluidity at low temperature (Nedwell, 1999) but provides an unrealistically high abundance of labile substrate to microorganisms when respiration is measured at a higher temperature. Soil microbial communities incubated at 10 °C may have become thermally adapted and able to maintain similar levels of metabolism at both 5 °C and 15 °C. There was no significant difference between the Q10 of soils incubated at an oscillating temperature between 5 °C and 15 °C and soils incubated at any of the constant temperatures (Figure 3). It seems that soils that had previously experienced time at 15 °C (constantly or oscillating between 5 °C and 15 °C) may have experienced the optimum temperature for microbial activity in the grassland soil where samples were collected and prevented any build-up of labile carbon, or thermal adaptation to colder temperatures. The temperature optimum of soil respiration is a reflection of long term physiological adaptation of soil microbial community to climate and environment (Rinnan et al., 2009; Liu et al., 2018).

### 5.2 Shifts in the soil microbial community structure in response to temperature regimes

An explanation for our observations regarding the legacy of prior incubation temperature on soil respiration is that the soil microbial community could have shifted in response to the temperatures that they were incubated at, as observed by Bradford et al., (2010). It is clear from global datasets of soil microbial communities that lower soil respiration at lower temperatures is indicative of the development of soil microbial communities with slower metabolic activities, such as fungi (Crowther et al., 2019), leading to the accumulation of organic carbon in fungal dominated ecosystems in colder climates.

Along with higher rates of soil respiration, we observed a shift away from a fungal dominated microbial community to one dominated more by gram-positive bacteria in soils incubated at a higher (or diurnally oscillating) temperature (Figure 6). This shift is consistent with observations made in the literature from experiments undertaken under warming conditions in both field and laboratory incubation experiments (Frey et al. 2008; Salazar et al. 2019). Our results therefore lend support to the general hypothesis that soils with a lower fungal-to-bacterial ratio have a lower potential to accumulate soil organic matter due to lower carbon use efficiency (Malik et al. 2016; Bonner et al. 2018). Greater dominance of fungi and gram negative bacteria have also been observed in soils with more total and labile carbon (Whitaker et al., 2014;Fanin et al., 2019). Therefore, the presence of greater total, and hot and cold water extractable organic carbon that we observed in soils incubated at lower temperatures (Figure 4) may have caused, or been the result of, shifts in the soil microbial community that raised the fungal-to-bacterial ratio and gram negative-to-gram positive bacteria ratio and may constitute an indirect mechanism by which soils respond to changes in temperature.

### 5.3 Depletion of soil organic matter at higher or oscillating incubation temperatures

Walker et al. (2018) identified a role for both substrate depletion and a permanent acceleration in microbial physiology that leads to faster respiration, growth, and turnover in warmed soils. Like Zogg et al. (1997), we observed differences in soil microbial community composition (Figure 6) between soils incubated at different temperatures that correspond with substrate depletion and soil respiration. Our laboratory incubation experiment revealed, at the higher incubation temperatures, elevated soil CO_2_ flux between week 5 and week 11, after which CO_2_ flux then decreased over time. This observation can most plausibly be explained by depletion of the labile soil organic carbon supply after 11 weeks. This explanation is consistent with the results from You et al. (2019), who observed a decrease in soil CO_2_ flux due to increasing temperature towards the end of a 35 day soil incubation study.

The lower concentration of CWEOC (Figure 4A) measured in soils incubated constantly at 15 °C or diurnally oscillated between 5 °C and 15 °C reveals that these soils have been depleted of dissolved organic carbon (DOC). This observation supports the results of Bertolet et al. (2018) who reported lower DOC under warmer temperatures in a 28 day incubation study. In another short term experiment, it was reported that increasing temperature reduced the DOC and microbial biomass carbon without any significant changes in soil organic matter or total C (You et al., 2019). In this study, we found higher DOC in soils incubated at 5 °C and 10 °C, compared to those incubated at 15 °C or oscillated between 5 °C and 15 °C. This observation indicates that a similar rate of substrate depletion occurred in soil incubated constantly at a high temperature and soils which oscillated between high and low temperatures. Our findings therefore imply that daily maximum temperature plays a more important role in soil organic matter (de)stabilisation than daily mean temperatures, complementing our interpretation of the CO_2_ flux measurements. The notion that the availability of low molecular weight compounds limits microbial intracellular respiration in our soils, and not the availability of a stoichiometric supply of nutrients, is supported by our results, which reveal greater N mineralisation in soils incubated at 15 °C or oscillated between 5 °C and 15 °C, compared with those incubated at 10 °C or 5 °C.

The lower C/N ratio in soils incubated at higher temperatures (Figure 4F) lends support to our interpretation that, at higher temperatures, readily available C is being depleted and N being mineralised. This evidence, coupled with the lack of leaching or plant uptake allowed in the experiment, indicates that the C left in the soil is more microbially processed (Bach et al., 2018) and more stable. The chemical fractionations of C and N in the soils incubated under oscillating temperature are most similar to the constant 15 °C incubation treatment, indicating that changes to C and N are dictated by maximum daily temperature rather than average daily temperature or minimum daily temperature. Soil microorganisms adapt to these changes in C and N availability to fulfil their energy and nutrient demands, thus causing shift in microbial community composition and physiology (Wan et al., 2014; Schnecker et al., 2015). However, shifts in community composition may also occur due to different groups of organisms outcompeting others for resources at the given temperature (Crowther et al., 2014).

### 5.4 Adaptation of the soil microbial community

In response to the stress associated with higher temperatures and substrate depletion, microorganisms are able to alter the composition of their cell walls to increase membrane stability (acclimatisation), but it is not possible, using fatty acid biomarkers, to distinguish between this phenomenon and a shift in the composition of the microbial community to one that comprises organisms with inherently more stable membranes (Frostegård et al., 2011). Commonly used microbial stress indicators include changes to the ratio of cyclopropyl fatty acids to their cis mono-unsaturated precursors, the ratio of gram-negative/gram-positive bacteria, cis/trans ratio, and iso/anteiso branching ratio (Kaur et al. 2005; Feng and Simpson 2009; Ruess and Chamberlain 2010; Sizmur et al. 2011; Willers et al. 2015; Bai et al. 2017). In our study, we found that higher incubation temperatures resulted in (i) a lower ratio of gram-negative/gram-positive bacteria biomarkers, (ii) a higher ratio of cy17:0c/16:1ω7c, (iii) a higher ratio of cis/trans ratio isomerization, and (iv) a higher iso/anteiso branching.

Both gram-negative/gram-positive ratio and the ratio of cyclopropyl fatty acids to their cis mono-unsaturated fatty acids precursors (cy17:0c/16:1ω7c in this study) are known indicators of temperature-induced nutrient depletions (Bai et al., 2017). Changes in cis/trans ratio isomerisation, and iso/anteiso branching have also been used to explain bacterial physiological adaptations (Ruess and Chamberlain, 2010) under stress conditions. The combination of these indicators could also represent microbial adaptation to sub-optimal temperature changes (Siliakus et al., 2017). Stress indicators were similar, although slightly lower, in soils oscillated between 5 °C and 15 °C, compared to soils incubated constantly at 15 °C, indicating that the changes to the phospholipid bilayer may be more associated with temperature than substrate depletion. Microbial adaptation, in the form of changing membrane composition, helps the community to combat environmental change (Feng and Simpson, 2009; De Maayer et al., 2014; Siliakus et al., 2017). Significantly lower ratios of cyclopropyl fatty acids to their cis mono-unsaturated fatty acid precursors (cy17:0c/16:1ω7c) and the iso/anteiso ratio in soils incubated under oscillating temperature, compared to those soils incubated constantly at 15 °C, reveals that temperature effects on the microbial community are lower in diurnally oscillating soils, compared to soils incubated at constant temperature. This observation could be because the oscillating treatments makes the best use of the diversity of the microbial community in the oscillating treatment, since different species may be capable of occupying different ‘temperature niches’ in a fluctuating environment (Upton et al., 1990). Our results thus imply that soil microbial communities incubated in fluctuating environments are less sensitive to change, compared to those incubated under constant conditions (Hawkes and Keitt, 2015).

## 6 Conclusions

We demonstrate that the daily maximum temperature a soil is exposed to has an important impact on soil microbial community composition, the rate and temperature sensitivity of soil respiration, and the depletion of soil organic matter in a temperate grassland soil. Our findings suggest that the daily maximum temperature mediates the production of extracellular enzymes which are capable of depolymerising macromolecules at lower temperatures overnight in sufficient quantity to maintain intracellular respiration. Microbial communities undergo changes to their composition and physiology when incubated under different temperature regimes. Lower temperatures shift the population towards a fungal dominated soil microbial community. Higher temperatures shift the population towards a bacterial dominated community with more gram-negative bacteria and greater membrane stability, due to thermal adaptation and in response to the stress associated with substrate depletion. The microbial communities of soils oscillated diurnally between 5 °C and daily 15 °C were similar to those maintained constantly at 15 °C. However, incubating soils at oscillating temperatures allows communities to exploit several different ‘temperature niches’. This knowledge is critical to advance new soil biogeochemical models that predict the impact of environmental change on soil respiration because asymmetric warming and a dampening of the diurnal temperature range is known to be occurring. It is recommended that the short-term impact of daily maximum and daily minimum temperatures on extracellular and intracellular enzyme activities, and the long-term impact of climate shifts on microbial community composition and physiology are incorporated into the next generation of soil carbon models. Soil respiration assays performed on soils pre-incubated at realistic temperatures that are representative of the daily maximum, daily average, and daily and minimum temperatures of the site from which the soils are samples would generate useful assessments of the likely temperature sensitivity (Q10) of soil respiration to future environmental change.

## Acknowledgement

This work was funded through the Commonwealth Scholarship granted to Adekanmbi A. A. to pursue a PhD in Soil Science.

## References

Abramoff, R., Xu, X., Hartman, M., O’Brien, S., Feng, W., Davidson, E., Finzi, A., Moorhead, D., Schimel, J., Torn, M., Mayes, M., 2017. The Millennial model: in search of measurable pools and transformations for modeling soil carbon in the new century. Biogeochemistry 137, 51–71. https://doi.org/10.1007/s10533-017-0409-7

Adekanmbi, A.A., Shaw, L.J., Sizmur, T., 2020. Effect of Sieving on Ex Situ Soil Respiration of Soils from Three Land Use Types. Journal of Soil Science and Plant Nutrition. https://doi.org/10.1007/s42729-020-00177-2

Akinremi, O.O., McGinn, S.M., McLean, H.D.J., 1999. Effects of soil temperature and moisture on soil respiration in barley and fallow plots. Canadian Journal of Soil Science 79, 5–13. https://doi.org/10.4141/S98-023

Alexander, L. V., Zhang, X., Peterson, T.C., Caesar, J., Gleason, B., Klein Tank, A.M.G., Haylock, M., Collins, D., Trewin, B., Rahimzadeh, F., Tagipour, A., Rupa Kumar, K., Revadekar, J., Griffiths, G., Vincent, L., Stephenson, D.B., Burn, J., Aguilar, E., Brunet, M., Taylor, M., New, M., Zhai, P., Rusticucci, M., Vazquez-Aguirre, J.L., 2006. Global observed changes in daily climate extremes of temperature and precipitation. Journal of Geophysical Research Atmospheres 111, 1–22. https://doi.org/10.1029/2005JD006290

Allison, S.D., 2014. Modeling adaptation of carbon use efficiency in microbial communities. Frontiers in Microbiology 5, 1–9. https://doi.org/10.3389/fmicb.2014.00571

Antwis, R.E., Griffiths, S.M., Harrison, X.A., Aranega-bou, P., Arce, A., Bettridge, A.S., Brailsford, F.L. Menezes, A. De, Devaynes, A., Forbes, K.M., Fry, E.L., Goodhead, I., Haskell, E., Heys, C., James, C., Johnston, S.R., Lewis, G.R., Lewis, Z., Macey, M.C., Mccarthy, A., Mcdonald, J.E., Mejia-florez, N.L., Brien, D.O., Pautasso, M., Reid, W.D.K., Robinson, H.A., Wilson, K., Sutherland, W.J., 2017. Fifty important research questions in microbial ecology 1–11. https://doi.org/10.1093/femsec/fix044

Bach, E.M., Williams, R.J., Hargreaves, S.K., Yang, F., Hofmockel, K.S., 2018. Greatest soil microbial diversity found in micro-habitats. Soil Biology and Biochemistry 118, 217–226. https://doi.org/10.1016/j.soilbio.2017.12.018

Bai, Z., Ma, Q., Wu, X., Zhang, Y., Yu, W., 2017. Temperature sensitivity of a PLFA-distinguishable microbial community differs between varying and constant temperature regimes. Geoderma 308, 54–59. https://doi.org/10.1016/j.geoderma.2017.08.026

Bardgett, R.D., Freeman, C., Ostle, N.J., 2008. Microbial contributions to climate change through carbon cycle feedbacks. ISME Journal 2, 805–814. https://doi.org/10.1038/ismej.2008.58

Bell, T.H., Klironomos, J.N., Henry, H.A.L., 2010. Seasonal Responses of Extracellular Enzyme Activity and Microbial Biomass to Warming and Nitrogen Addition. Soil Science Society of America Journal 74, 820. https://doi.org/10.2136/sssaj2009.0036

Bertolet, B.L., Corman, J.R., Casson, N.J., Sebestyen, S.D., Kolka, R.K., Stanley, E.H., 2018. Influence of soil temperature and moisture on the dissolved carbon, nitrogen, and phosphorus in organic matter entering lake ecosystems. Biogeochemistry 139, 293–305. https://doi.org/10.1007/s10533-018-0469-3

Bonner, M.T.L., Shoo, L.P., Brackin, R., Schmidt, S., 2018. Geoderma Relationship between microbial composition and substrate use e ffi ciency in a tropical soil. Geoderma 315, 96–103. https://doi.org/10.1016/j.geoderma.2017.11.026

Bradford, M.A., Watts, B.W., Davies, C.A., 2010. Thermal adaptation of heterotrophic soil respiration in laboratory microcosms. Global Change Biology 16, 1576–1588. https://doi.org/10.1111/j.1365-2486.2009.02040.x

Braganza, K., Karoly, D.J., Arblaster, J.M., 2004. Diurnal temperature range as an index of global climate change during the twentieth century. Geophysical Research Letters 31, 2–5. https://doi.org/10.1029/2004GL019998

Chang, W., Whyte, L., Ghoshal, S., 2011. Comparison of the effects of variable site temperatures and constant incubation temperatures on the biodegradation of petroleum hydrocarbons in pilot-scale experiments with field-aged contaminated soils from a cold regions site. Chemosphere 82, 872–878. https://doi.org/10.1016/j.chemosphere.2010.10.072

Chen, J.M., Huang, S.E., Ju, W., Gaumont-Guay, D., Black, T.A., 2009. Daily heterotrophic respiration model considering the diurnal temperature variability in the soil G01022. Journal of Geophysical Research: Biogeosciences 114, 1–11. https://doi.org/10.1029/2008JG000834

Conant, R.T., Steinweg, J.M., Haddix, M.L., Paul, E.A., Plante, A.F., Six, J., Six, J., 2008. Experimental Warming Shows That Decomposition Temperature Sensitivity Increases with Soil Organic Matter Recalcitrance Published by?: Wiley on behalf of the Ecological Society of America Stable URL?: http://www.jstor.com/stable/27650775 REFERENCES Linked 89, 2384–2391.

Crowther, T.W., Maynard, D.S., Crowther, T.R., Peccia, J., Smith, J.R., Bradford, M.A., 2014. Untangling the fungal niche: The trait-based approach. Frontiers in Microbiology 5, 1–12. https://doi.org/10.3389/fmicb.2014.00579

Crowther, T.W., van den Hoogen, J., Wan, J., Mayes, M.A., Keiser, A.D., Mo, L., Averill, C., Maynard, D.S., 2019. The global soil community and its influence on biogeochemistry. Science (New York, N.Y.) 365. https://doi.org/10.1126/science.aav0550

Curiel Yuste, J., Janssens, I.A., Carrara, A., Ceulemans, R., 2004. Annual Q10 of soil respiration reflects plant phenological patterns as well as temperature sensitivity. Global Change Biology 10, 161–169. https://doi.org/10.1111/j.1529-8817.2003.00727.x

Davidson, E.A., Janssens, I.A., 2006. Temperature sensitivity of soil carbon decomposition and feedbacks to climate change. Nature 440, 165–173. https://doi.org/10.1038/nature04514

Davidson, E.A., Janssens, I.A., Lou, Y., 2006. On the variability of respiration in terrestrial ecosystems: Moving beyond Q10. Global Change Biology 12, 154–164. https://doi.org/10.1111/j.1365-2486.2005.01065.x

De Maayer, P., Anderson, D., Cary, C., Cowan, D.A., 2014. Some like it cold: Understanding the survival strategies of psychrophiles. EMBO Reports 15, 508–517. https://doi.org/10.1002/embr.201338170

Dondini, M., Richards, M., Pogson, M., Jones, E.O., Rowe, R.L., Keith, A.M., McNamara, N.P., Smith, J.U., Smith, P., 2016. Evaluation of the ECOSSE model for simulating soil organic carbon under Miscanthus and short rotation coppice-willow crops in Britain. GCB Bioenergy 8, 790–804. https://doi.org/10.1111/gcbb.12286

Dutta, H., Dutta, A., 2016. The microbial aspect of climate change. Energy, Ecology and Environment 1, 209–232. https://doi.org/10.1007/s40974-016-0034-7

Fang, C., Smith, P., Moncrieff, J.B., Smith, J.U., 2005. Similar response of labile and resistant soil organic matter pools to changes in temperature 57–59. https://doi.org/10.1130/G20750

Fanin, N., Kardol, P., Farrell, M., Nilsson, M., Gundale, M.J., Wardle, D.A., 2019. The ratio of Gram-positive to Gram-negative bacterial PLFA markers as an indicator of carbon availability in organic soils. Soil Biology and Biochemistry 128, 111–114. https://doi.org/10.1016/j.soilbio.2018.10.010

Feng, X., Simpson, M.J., 2009. Temperature and substrate controls on microbial phospholipid fatty acid composition during incubation of grassland soils contrasting in organic matter quality. Soil Biology and Biochemistry 41, 804–812. https://doi.org/10.1016/j.soilbio.2009.01.020

Foereid, B., Ward, D.S., Mahowald, N., Paterson, E., Lehmann, J., 2014. The sensitivity of carbon turnover in the Community Land Model to modified assumptions about soil processes. Earth System Dynamics 5, 211–221. https://doi.org/10.5194/esd-5-211-2014

Frey, S.D., Drijber, R., Smith, H., Melillo, J., 2008. Microbial biomass, functional capacity, and community structure after 12 years of soil warming. Soil Biology and Biochemistry 40, 2904–2907. https://doi.org/10.1016/j.soilbio.2008.07.020

Frostegård, Å., Tunlid, A., Bååth, E., 2011. Use and misuse of PLFA measurements in soils. Soil Biology and Biochemistry 43, 1621–1625. https://doi.org/10.1016/j.soilbio.2010.11.021

Ghani, A., Dexter, M., Perrott, K.W., 2003. Hot-Water Extractable Carbon in Soils?: A Sensitive Measurement for Determining Impacts of Fertilisation, Grazing and Cultivation Hot-water extractable carbon in soils?: a sensitive measurement for determining impacts of fertilisation, grazing and culti 717. https://doi.org/10.1016/S0038-0717(03)00186-X

Gilhespy, S.L., Anthony, S., Cardenas, L., Chadwick, D., del Prado, A., Li, C., Misselbrook, T., Rees, R.M., Salas, W., Sanz-Cobena, A., Smith, P., Tilston, E.L., Topp, C.F.E., Vetter, S., Yeluripati, J.B., 2014. First 20 years of DNDC (DeNitrification DeComposition): Model evolution. Ecological Modelling 292, 51–62. https://doi.org/10.1016/j.ecolmodel.2014.09.004

Graf, A., Weihermüller, L., Huisman, J.A., Herbst, M., Bauer, J., Vereecken, H., 2008. Measurement depth effects on the apparent temperature sensitivity of soil respiration in field studies. Biogeosciences 5, 1175–1188. https://doi.org/10.5194/bg-5-1175-2008

Gritsch, C., Zimmermann, M., Zechmeister-Boltenstern, S., 2015. Interdependencies between temperature and moisture sensitivities of CO2emissions in European land ecosystems. Biogeosciences 12, 5981–5993. https://doi.org/10.5194/bg-12-5981-2015

Hansen, J., Sato, M., Ruedy, R., 1995. Long-term changes of the diurnal temperature cycle: implications about mechanisms of global climate change. Atmospheric Research 37, 175–209. https://doi.org/10.1016/0169-8095(94)00077-Q

Harden, J.W., Hugelius, G., Ahlström, A., Blankinship, J.C., Bond-Lamberty, B., Lawrence, C.R., Loisel, J., Malhotra, A., Jackson, R.B., Ogle, S., Phillips, C., Ryals, R., Todd-Brown, K., Vargas, R., Vergara, S.E., Cotrufo, M.F., Keiluweit, M., Heckman, K.A., Crow, S.E., Silver, W.L., DeLonge, M., Nave, L.E., 2018. Networking our science to characterize the state, vulnerabilities, and management opportunities of soil organic matter. Global Change Biology 24, e705–e718. https://doi.org/10.1111/gcb.13896

Hawkes, C. V., Keitt, T.H., 2015. Resilience vs. historical contingency in microbial responses to environmental change. Ecology Letters 18, 612–625. https://doi.org/10.1111/ele.12451

Jan, M.T., Roberts, P., Tonheim, S.K., Jones, D.L., 2009. Protein breakdown represents a major bottleneck in nitrogen cycling in grassland soils. Soil Biology and Biochemistry 41, 2272–2282. https://doi.org/10.1016/j.soilbio.2009.08.013

Karhu, K., Auffret, M.D., Dungait, J.A.J., Hopkins, D.W., Prosser, J.I., Singh, B.K., Subke, J.A., Wookey, P.A., Agren, G.I., Sebastià, M.T., Gouriveau, F., Bergkvist, G., Meir, P., Nottingham, A.T., Salinas, N., Hartley, I.P., 2014. Temperature sensitivity of soil respiration rates enhanced by microbial community response. Nature 513, 81–84. https://doi.org/10.1038/nature13604

Kaur, A.A., Chaudhary, A., Choudhary, R., Kaushik, R., Kaur, A.A., Choudhary, R., Kaushik, R., 2005. Phospholipid fatty acid – A bioindicator of environment monitoring and assessment in soil ecosystem. October 89, 1103–1112. https://doi.org/10.2307/24110962

Kirschbaum, M.U.F.F., 1995. The temperature dependence of soil organic matter decomposition, and the effect of global warming on soil organic C storage. Soil Biology and Biochemistry 27, 753–760. https://doi.org/10.1016/0038-0717(94)00242-S

Li, J., He, N., Wei, X., Gao, Y., Zuo, Y., 2015. Changes in temperature sensitivity and activation energy of soil organic matter decomposition in different Qinghai-Tibet Plateau grasslands. PLoS ONE 10, 1–14. https://doi.org/10.1371/journal.pone.0132795

Lin, J., Zhu, B., Cheng, W., 2015. Decadally cycling soil carbon is more sensitive to warming than faster-cycling soil carbon. Global Change Biology 21, 4602–4612. https://doi.org/10.1111/gcb.13071

Liu, Y., He, N., Wen, X., Xu, L., Sun, X., Yu, G., Liang, L., Schipper, L.A., 2018. The optimum temperature of soil microbial respiration: Patterns and controls. Soil Biology and Biochemistry 121, 35–42. https://doi.org/10.1016/j.soilbio.2018.02.019

Malik, A.A., Chowdhury, S., Schlager, V., Oliver, A., Puissant, J., Vazquez, P.G.M., Jehmlich, N., von Bergen, M., Griffiths, R.I., Gleixner, G., 2016. Soil fungal: Bacterial ratios are linked to altered carbon cycling. Frontiers in Microbiology 7, 1–11. https://doi.org/10.3389/fmicb.2016.01247

Melillo, J.M., Frey, S.D., DeAngelis, K.M., Werner, W.J., Bernard, M.J., Bowles, F.P., Pold, G., Knorr, M.A., Grandy, A.S., 2017. Long-term pattern and magnitude of soil carbon feedback to the climate system in a warming world. Science 358, 101–105. https://doi.org/10.1126/science.aan2874

Meng, C., Tian, D., Zeng, H., Li, Z., Chen, H.Y.H., Niu, S., 2020. Global meta-analysis on the responses of soil extracellular enzyme activities to warming. Science of the Total Environment 705, 135992. https://doi.org/10.1016/j.scitotenv.2019.135992

Meyer, N., Welp, G., Amelung, W., 2019. Effect of sieving and sample storage on soil respiration and its temperature sensitivity (Q10) in mineral soils from Germany. Biology and Fertility of Soils 55, 833. https://doi.org/10.1007/s00374-019-01380-9

Meyer, N., Welp, G., Amelung, W., 2018. The Temperature Sensitivity (Q10) of Soil Respiration: Controlling Factors and Spatial Prediction at Regional Scale Based on Environmental Soil Classes. Global Biogeochemical Cycles 32, 306–323. https://doi.org/10.1002/2017GB005644

Mitra, B., Miao, G., Minick, K., McNulty, S.G., Sun, G., Gavazzi, M., King, J.S., Noormets, A., 2019. Disentangling the Effects of Temperature, Moisture, and Substrate Availability on Soil CO2 Efflux. Journal of Geophysical Research: Biogeosciences 124, 2060–2075. https://doi.org/10.1029/2019JG005148

Nedwell, D.B., 1999. Effect of low temperature on microbial growth: Lowered affinity for substrates limits growth at low temperature. FEMS Microbiology Ecology 30, 101–111. https://doi.org/10.1016/S0168-6496(99)00030-6

Oksanen, J., 2017. Vegan: ecological diversity. R Package Version 2.4-4 1, 11.

Parton, W.J., 1996. The CENTURY model, in: Powlson, D.S., Smith, P., Smith, J.U. (Eds.), Evaluation of Soil Organic Matter Models. Springer, Berlin, Heidelberg, pp. 283–291.

Pavelka, M., Acosta, M., Marek, M. V., Kutsch, W., Janous, D., 2007. Dependence of the Q10values on the depth of the soil temperature measuring point. Plant and Soil 292, 171–179. https://doi.org/10.1007/s11104-007-9213-9

Pold, G., Grandy, A.S., Melillo, J.M., DeAngelis, K.M., 2017. Changes in substrate availability drive carbon cycle response to chronic warming. Soil Biology and Biochemistry 110, 68–78. https://doi.org/10.1016/j.soilbio.2017.03.002

Potter, C.S., Randerson James T. Field, Christopher B. Matson, P.A., Vitousek, P.M., Mooney, H.A., Klooster, S.A., 1993. Terrestrial ecosystem production: A process model based on global satellite and surface data. Global Biogeochemical Cycles 7, 811–841. https://doi.org/10.1029/93GB02725

Quideau, S.A., McIntosh, A.C.S., Norris, C.E., Lloret, E., Swallow, M.J.B., Hannam, K., 2016. Extraction and analysis of microbial Phospholipid fatty acids in soils. Journal of Visualized Experiments 2016, 1–9. https://doi.org/10.3791/54360

Raich, J.W., Rastetter, E.B., Melillo, J.M., Kicklighter, D.W., Steudler, A.P., Peterson, B.J., Schloss, A.L., Moore, I.B., Vorosmarty, C.J., 1991. Potential net primary productivity in South America: application of a global model. Ecological Applications 1, 399–429. https://doi.org/10.2307/1941899

Rinnan, R., Rousk, J., Yergeau, E., Kowalchuk, G.A., Bååth, E., 2009. Temperature adaptation of soil bacterial communities along an Antarctic climate gradient: Predicting responses to climate warming. Global Change Biology 15, 2615–2625. https://doi.org/10.1111/j.1365-2486.2009.01959.x

Rodriguez-Verdugo, A., Vulin, C., Ackermann, M., 2019. The rate of environmental fluctuations shapes ecological dynamics in a two-species microbial system. Ecology letters 22, 838–846. https://doi.org/10.1111/ele.13241

Ross, D.J., Täte, K.R., 1993. Microbial C and N, and respiratory activity, in litter and soil of a southern beech (Nothofagus) forest: Distribution and properties. Soil Biology and Biochemistry 25, 477–483. https://doi.org/10.1016/0038-0717(93)90073-K

Ruess, L., Chamberlain, P.M., 2010. The fat that matters: Soil food web analysis using fatty acids and their carbon stable isotope signature. Soil Biology and Biochemistry 42, 1898–1910. https://doi.org/10.1016/j.soilbio.2010.07.020

Salazar, A., Rousk, K., Jónsdóttir, I.S., Bellenger, J., Andrésson, Ó.S., 2019. Faster nitrogen cycling and more fungal and root biomass in cold ecosystems under experimental warming: a meta-analysis. Ecology 0, 1–13. https://doi.org/10.1002/ecy.2938

Schimel, J.P., Weintraub, M.N., 2003. The implications of exoenzyme activity on microbial carbon and nitrogen limitation in soil: A theoretical model. Soil Biology and Biochemistry 35, 549–563. https://doi.org/10.1016/S0038-0717(03)00015-4

Schmidt, M.W.I., Torn, M.S., Abiven, S., Dittmar, T., Guggenberger, G., Janssens, I.A., Kleber, M. Kögel-Knabner, Ingrid Lehmann, J., Manning, D.A.C., Nannipieri, P., Rasse, D.P., Weiner, S., Trumbore, S.E., 2011. Persistence of soil organic matter as an ecosystem property. Nature 478, 49–56.

Schnecker, J., Wild, B., Takriti, M., Eloy Alves, R.J., Gentsch, N., Gittel, A., Hofer, A., Klaus, K., Knoltsch, A., Lashchinskiy, N., Mikutta, R., Richter, A., 2015. Microbial community composition shapes enzyme patterns in topsoil and subsoil horizons along a latitudinal transect in Western Siberia. Soil Biology and Biochemistry 83, 106–115. https://doi.org/10.1016/j.soilbio.2015.01.016

Shurpali, N.J., Rannik, Ü., Jokinen, S., Lind, S., Biasi, C., Mammarella, I., Peltola, O., Pihlatie, M., Hyvönen, N., Räty, M., Haapanala, S., Zahniser, M., Virkajärvi, P., Vesala, T., Martikainen, P.J., 2016. Neglecting diurnal variations leads to uncertainties in terrestrial nitrous oxide emissions. Nature Publishing Group 1–10. https://doi.org/10.1038/srep25739

Siliakus, M.F., van der Oost, J., Kengen, S.W.M., 2017. Adaptations of archaeal and bacterial membranes to variations in temperature, pH and pressure. Extremophiles 21, 651–670. https://doi.org/10.1007/s00792-017-0939-x

Sizmur, T., Tilston, E.L., Charnock, J., Palumbo-Roe, B., Watts, M.J., Hodson, M.E., 2011. Impacts of epigeic, anecic and endogeic earthworms on metal and metalloid mobility and availability. Journal of Environmental Monitoring 13, 266–273. https://doi.org/10.1039/c0em00519c

Smith, K.A., Ball, T., Conen, F., Dobbie, K.E., Massheder, J., Rey, A., 2018. Exchange of greenhouse gases between soil and atmosphere: interactions of soil physical factors and biological processes. European Journal of Soil Science 69, 10–20. https://doi.org/10.1111/ejss.12539

Subke, J.A., Bahn, M., 2010. On the “temperature sensitivity” of soil respiration: Can we use the immeasurable to predict the unknown? Soil Biology and Biochemistry 42, 1653–1656. https://doi.org/10.1016/j.soilbio.2010.05.026

Thiessen, S., Gleixner, G., Wutzler, T., Reichstein, M., 2013. Both priming and temperature sensitivity of soil organic matter decomposition depend on microbial biomass - An incubation study. Soil Biology and Biochemistry 57, 739–748. https://doi.org/10.1016/j.soilbio.2012.10.029

Todd-Brown, K., Hopkins, F.M., Kivlin, S.N., Talbot, J.M., Allison, S.D., 2012. A framework for representing microbial decomposition in coupled climate models. Biogeochemistry. https://doi.org/10.1007/s10533-011-9635-6

Upton, A.C., Nedwell, D.B., Wynn-Williams, D.D., 1990. The selection of microbial communities by constant or fluctuating temperatures. FEMS Microbiology Letters 74, 243–252. https://doi.org/10.1111/j.1574-6968.1990.tb04070.x

von Lützow, M., Kögel-Knabner, I., 2009. Temperature sensitivity of soil organic matter decomposition-what do we know? Biology and Fertility of Soils 46, 1–15. https://doi.org/10.1007/s00374-009-0413-8

Walker, T.W.N., Kaiser, C., Strasser, F., Herbold, C.W., Leblans, N.I.W., Woebken, D., Janssens, I.A., Sigurdsson, B.D., Richter, A., 2018. Microbial temperature sensitivity and biomass change explain soil carbon loss with warming. Nature Climate Change 8, 885–889. https://doi.org/10.1038/s41558-018-0259-x

Whitaker, J., Ostle, N., Nottingham, A.T., Ccahuana, A., Salinas, N., Bardgett, R.D., Meir, P., Mcnamara, N.P., 2014. Microbial community composition explains soil respiration responses to changing carbon inputs along an Andes-to-Amazon elevation gradient. Journal of Ecology 102, 1058–1071. https://doi.org/10.1111/1365-2745.12247

Wieder, W.R., Bonan, G.B., Steven, D.A., 2013. Global soil carbon projections are improved by modelling microbial processes. Nature Climate Change 3, 909–912.

Willers, C., Jansen van Rensburg, P.J., Claassens, S., 2015. Phospholipid fatty acid profiling of microbial communities-a review of interpretations and recent applications. Journal of Applied Microbiology 119, 1207–1218. https://doi.org/10.1111/jam.12902

Winkler, J.P., Cherry, R.S., Schlesinger, W.H., 1996. THE Q,,, RELATIONSHIP OF MICROBIAL RESPIRATION IN A TEMPERATE FOREST SOIL A “ bey?; R?;’ 28, 1067–1072.

Wu, Y., Yu, X., Wang, H., Ding, N., Xu, J., 2010. Does history matter? Temperature effects on soil microbial biomass and community structure based on the phospholipid fatty acid (PLFA) analysis. Journal of Soils and Sediments 10, 223–230. https://doi.org/10.1007/s11368-009-0118-5

Xu, J., Wei, Q., Yang, S., Wang, Y., Lv, Y., 2016. Diurnal pattern of nitrous oxide emissions from soils under different vertical moisture distribution conditions. Chilean Journal of Agricultural Research 76, 48–92. https://doi.org/10.4067/S0718-58392016000100012

Yang, K., He, R., Yang, W., Li, Z., Zhuang, L., Wu, F., Tan, B., Liu, Y., Zhang, L., Tu, L., Xu, Z., 2017. Temperature response of soil carbon decomposition depends strongly on forest management practice and soil layer on the eastern Tibetan Plateau. Scientific Reports 7, 1–8. https://doi.org/10.1038/s41598-017-05141-2

You, G., Zhang, Z., Zhang, R., 2019. Temperature adaptability of soil respiration in short-term incubation experiments. Journal of Soils and Sediments 19, 557–565. https://doi.org/10.1007/s11368-018-2059-3

Zang, H., Blagodatskaya, E., Wen, Y., Shi, L., Cheng, F., Chen, H., Zhao, B., Zhang, F., Fan, M., Kuzyakov, Y., 2020. Temperature sensitivity of soil organic matter mineralization decreases with long-term N fertilization: Evidence from four Q10 estimation approaches. Land Degradation and Development 31, 683–693. https://doi.org/10.1002/ldr.3496

Zhou, Z., Xu, M., Kang, F., Sun, O.J., 2015. Maximum temperature accounts for annual soil CO2 efflux in temperate forests of Northern China. Scientific Reports 5, 1–10. https://doi.org/10.1038/srep12142

Zogg, G.P., Zak, D.R., Ringelberg, D.B., White, D.C., MacDonald, N.W., Pregitzer, K.S., 1997. Compositional and Functional Shifts in Microbial Communities Due to Soil Warming. Soil Science Society of America Journal 61, 475. https://doi.org/10.2136/sssaj1997.03615995006100020015x

