## Supplementary material for "Legacy effect of constant and diurnally oscillating temperatures on soil respiration and microbial community structure"

**This supplementary material contains:**

- **A description of the laboratory procedures used for characterisation of soil samples prior to incubations**
- **Figure S-1: Effects incubation temperature on soil $CO_2$ released from soil when $CO_2$ flux was measured at 5 °C, 10 °C, and 15 °C**
- **Figure S-2: Effects measurement temperature on soil $CO_2$ released from soil previously incubated at 5 °C, 10 °C, 15 °C, or oscillating between 5 °C and 15 °C**
- **Table S-1 The designation of individual PLFA biomarkers to microbial community groups**
- **Supplementary References**

**A description of the laboratory procedures used for characterisation of soil samples prior to incubations**

Soil characteristics were measured using standard laboratory methods. The particle size distribution of soils was determined using a Malvern Mastersizer3000 Laser Granulometer after dispersing the soil in a solution containing 3.3% sodium hexametaphosphate + 0.7% sodium carbonate. The data was converted from % volume to % mass as described in(Yang et al., 2015) . Soil pH was determined by shaking soil samples with deionised water (1:10 mass/volume ratio) for 30 min and leaving the mixture to stand for 2 min before pH was measured using a digital type DMP-2 mV/pH meter (Thermo Orion). Total N and C concentrations were determined using C/N Elemental Analyser (Thermo Flash 2000 EA). The C/N ratio was then calculated from total C and N. Nitrate and ammonia were extracted in 1M KCl and then analysed using a Continuous Flow Analyzer (San++ Automated Wet Chemistry Analyzer - SKALAR). Moisture content and loss on ignition were determined by weight loss at 105 °C and 500 °C, respectively. Soil water holding capacity (WHC) was determined using saturation and drain method by submerging a 30 g air-dried sample in a plastic cylinder with a mesh bottom in water for 12 h to ensure complete saturation and then allowing the water to drain for another 12 h. The drained soil was then oven-dried at 105.

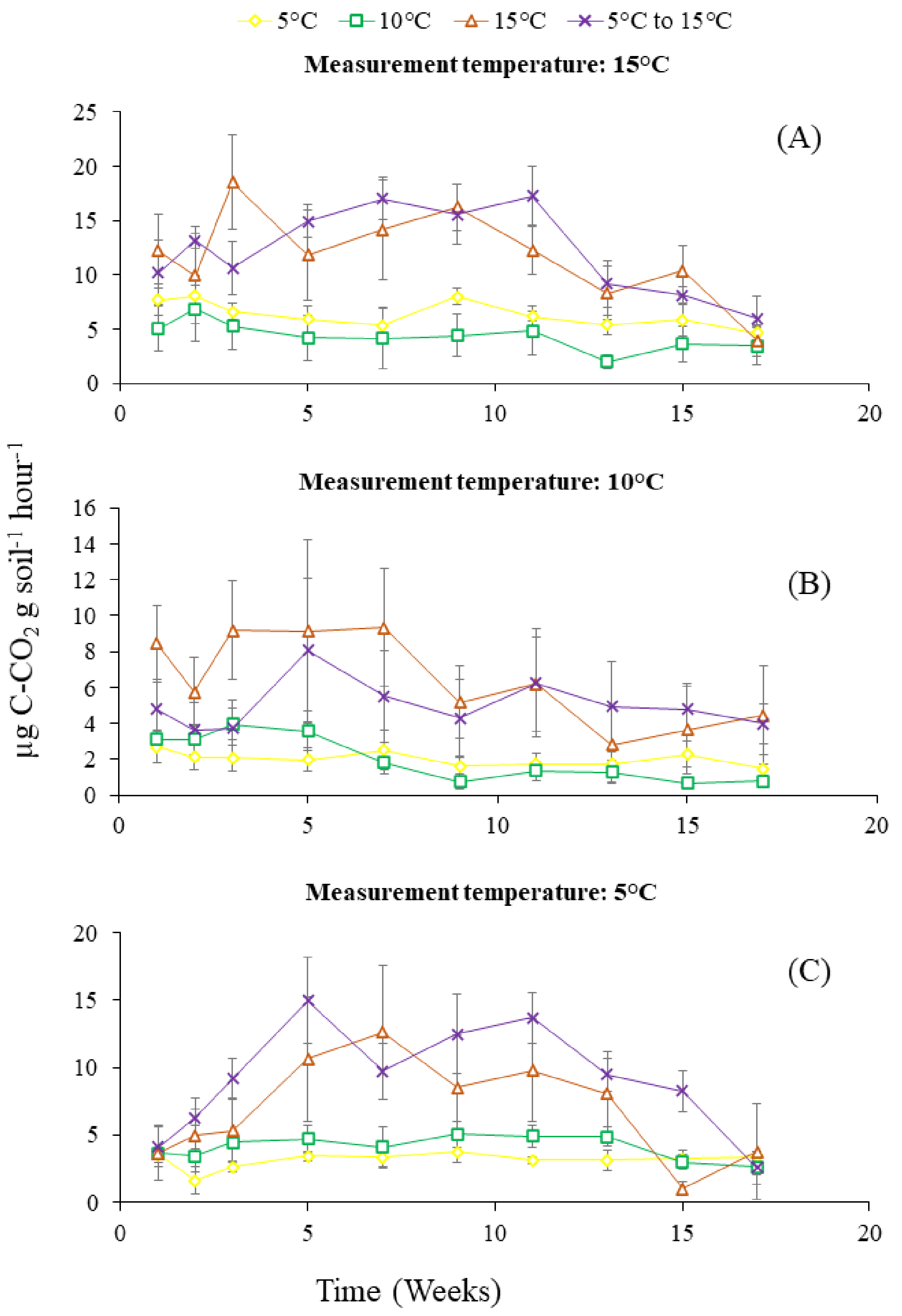


**Figure S-1: Effects of incubation temperature on soil $CO_2$ released from soil when $CO_2$ flux was measured at 5 °C, 10 °C, and 15 °C.**

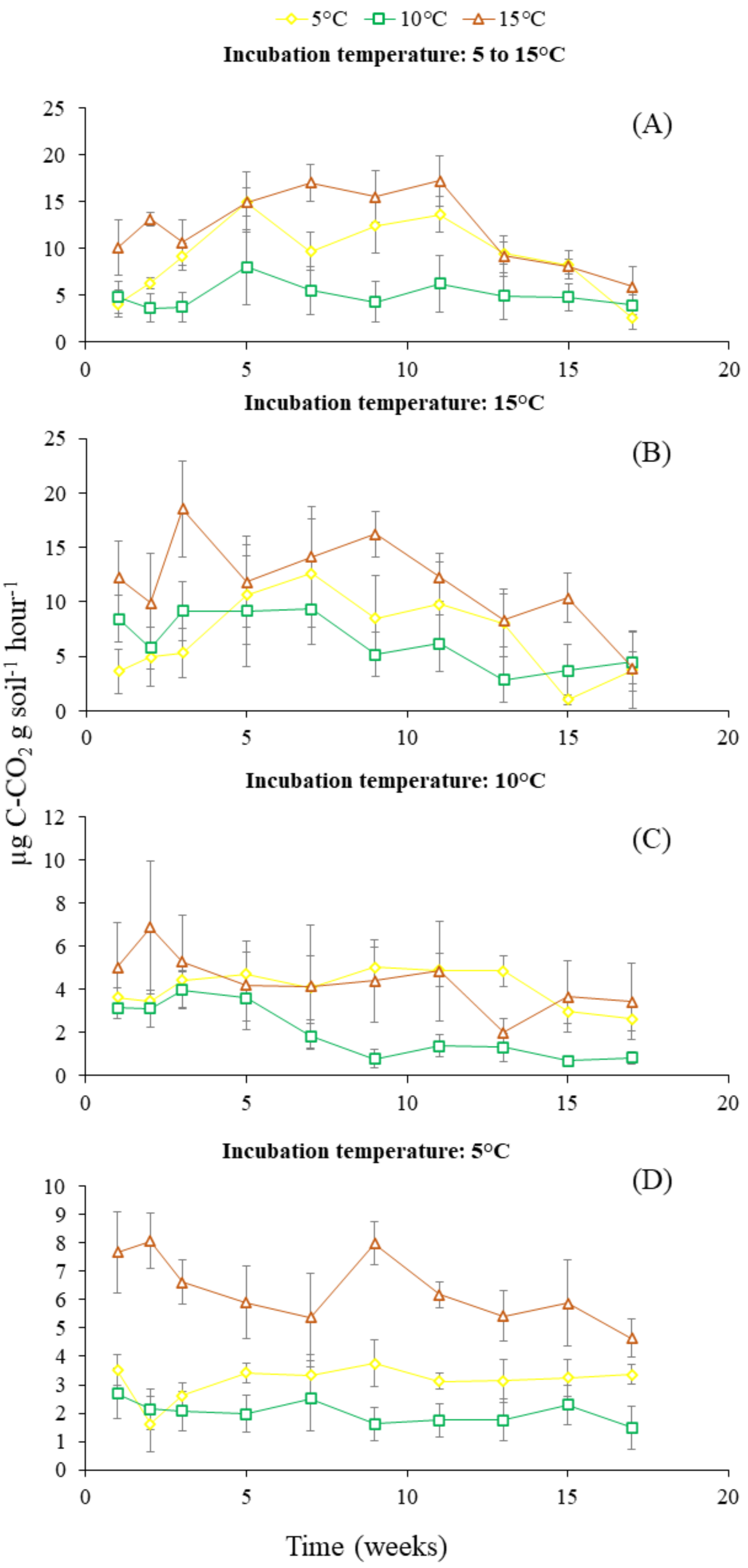


**Figure S-2: Effects of measurement temperature on soil $CO_2$ released from soil previously incubated at 5 °C, 10 °C, 15 °C, or oscillating between 5 °C and 15 °C.**

**Table S-1 The designation of individual PLFA biomarkers to microbial community groups**

| Group | Biomarker | Reference |
|---|---|---|
| Gram negative bacteria | α14:0 2OH, β14:0 3OH, cy17:0c, cy19:0c, 16:1ω7c, and 16:1w7t | Kaur et al., (2005), Willers et al., (2015) |
| Gram positive bacteria | i15:0, a15:0, i16:0, br17:0, and i17:0, | Kaur et al., (2005), Willers et al., (2015) |
| Bacteria | C15:0, C17:0, C20:0, α14:0 2OH, β14:0 3OH, cy17:0c, cy19:0c, 16:1ω7c, and 16:1w7t, i15:0, a15:0, i16:0, br17:0, and i17:0. | Bossio and Scow, (1998), Willers et al., (2015) |
| Fungi | 18:1ω9t, 18:2ω6t, and 18:2ω6,9c | Bossio and Scow, 1998), Kaur et al., (2005), |
| cis isomers | 16:1ω7c, 17:1c, cy17:0c, 18:3ω6c, 18:2ω6,9c, cy19:0c, 20:4ω6c, 20:5ω3c, 20:1ω9c, | Quideau et al., (2016) |
| trans isomers | C16:1w11t, 16:1w7t, 18:2ω6t, and 18:1ω9t | Quideau et al., (2016) |
| iso | i15:0, i16:0, and i17:0 | Quideau et al., (2016) |
| anteiso | a15:0 | Quideau et al., (2016) |

### Supplementary References


Bossio, D.A., Scow, K.M., 1998. Impacts of carbon and flooding on soil microbial communities: Phospholipid fatty acid profiles and substrate utilization patterns. Microbial Ecology 35, 265–278. doi:10.1007/s002489900082

Kaur, Amrit, Chaudhary, A., Kaur, Amarjeet, Choudhary, R., Kaushik, R., 2005. Phospholipid fatty acid - A bioindicator of environment monitoring and assessment in soil ecosystem. Current Science 89, 1103–1112.

Quideau, S.A., McIntosh, A.C.S., Norris, C.E., Lloret, E., Swallow, M.J.B., Hannam, K., 2016. Extraction and analysis of microbial Phospholipid fatty acids in soils. Journal of Visualized Experiments 2016, 1–9. doi:10.3791/54360

Willers, C., Jansen van Rensburg, P.J., Claassens, S., 2015. Phospholipid fatty acid profiling of microbial communities-a review of interpretations and recent applications. Journal of Applied Microbiology 119, 1207–1218. doi:10.1111/jam.12902

Yang, X., Zhang, Q., Li, X., Jia, X., Wei, X., Shao, M., 2015. Determination of Soil Texture by Laser Diffraction Method. Soil Science Society of America Journal 79, 1556. doi:10.2136/sssaj2015.04.0164
